# Mesoscale Ca^2+^ imaging reveals networks of Purkinje cell dendritic and somatic modulation, with divergent roles of activity versus correlation during behavior

**DOI:** 10.1101/2023.10.05.561090

**Authors:** M.L. Streng, R.E. Carter, B.W. Kottke, K. Togneri, E. Wasserman, V. Rajendran, S. B. Kodandaramaiah, E. Krook-Magnuson, T.J. Ebner

**Affiliations:** Department of Neuroscience, University of Minnesota, Minneapolis, MN, USA

## Abstract

A major challenge in cerebellar physiology is determining how the stereotypic, conserved circuitry of the cerebellar cortex, with its dominant parasagittal and transverse architectures, underlies its fundamental computations and contributions to behavior. To interrogate Purkinje cell dynamics at this parasagittal and transverse spatial scale, we implemented a novel approach for awake, chronic, wide-field Ca^2+^ imaging of the cerebellar cortex. We observe two functionally and spatially distinct Purkinje cell networks, reflecting their dendritic and somatic activities, respectively. Both dendritic and somatic networks exhibit bilateral, widespread activation during behavior, but with diverse patterns of spatial correlations occurring primarily along the parasagittal and transverse directions, consistent with the main geometry of the cerebellar cortex. Somatic network correlation dynamics are robustly modulated by prediction errors, and even reflect ultimate behavioral outcomes. These results provide a novel link between cerebellar structure and function, with the correlation dynamics of Purkinje cell activity a key feature.

## Introduction

The stereotypic and relatively simple circuitry of the cerebellar cortex, with its highly conserved nature throughout evolution has long intrigued neuroscientists. With three major classes of intrinsic neurons, two major classes of inputs, and the basic physiology defined, 50 years ago an understanding of cerebellar function seemed to be within reach^1–7^. Also, the organization of the circuitry, including the parasagittal olivo-cerebellar and cortico-nuclear projections, expression of molecular markers^8^ and the transverse organization of the parallel fibers, provides a natural and approachable spatial framework for the essential circuit computations. These features have led to the widely held hypothesis that the cerebellar cortex provides a uniform input-output transformation that is needed not only for motor function but a spectrum of non-motor functions^9^. Despite these longstanding views and fundamental observations, a unifying view of the computations performed by the cerebellar cortex that incorporates its dominant parasagittal and transverse architectures has proved elusive.

While new insights into the underlying molecular and cell properties have greatly enhanced our knowledge^10–15^, these spatial organizations are undoubtedly fundamental to cerebellar function. The parasagittal and transverse organizations suggest that requisite for uncovering the basic computations performed by the cerebellar cortex is their investigation at these corresponding spatial levels. The fractured somatotopy of the cerebellar cortex reinforces this need, as information about the body is distributed across multiple folia^16, 17^. Early efforts to investigate spatial processing in the cerebellar cortex relied on electrode arrays, providing important information on the synchrony and spatial activity of climbing fiber activity^18–20^. More recently, two-photon Ca^2+^ imaging of cerebellar neurons within a single folium has yielded new insights into the encoding of behavior by local neuronal populations^21–24^. However, recordings of cerebellar neuronal activity across multiple folia at relatively high spatial and temporal resolution are limited.

We investigated large-scale Purkinje cell activity across multiple folia at cellular resolution using a polymer window that allows for the resolution of Purkinje cell activity across multiple folia during chronic, awake behavior. In mice sparsely expressing GCaMP6s in Purkinje cells, we are able to record Purkinje cell activity across a large (∼13mm^2^) cortical region at the single cell level, including monitoring the dendrites and somata of hundreds of Purkinje cells. Blind source separation using spatial Independent Component Analysis (sICA) reveals two functional and spatially different networks of Purkinje cell activity, which correspond to either dendritic or somatic modulation. Both dendritic and somatic networks are highly engaged during spontaneous locomotion, cued reaching, and prediction errors, with widespread activation throughout the dorsal cerebellar cortex. Each behavior is characterized by distinct, modulated patterns of spatial correlations that differ for dendritic versus somatic networks. Occurring primarily along parasagittal and transverse directions, the changes in the correlation patterns strongly reflect the behavior. Somatic network dynamics are dramatically altered by prediction errors, whose processing is considered a hallmark of cerebellar function, and can even reflect ultimate behavioral outcomes. Together, these results provide new insights into the organization of Purkinje cell dendritic versus somatic information at the network level, highlight important differences between neuronal activation versus correlation, and outline Purkinje cell somatic network correlation dynamics as a potentially key feature of cerebellar processing.

## Results

### Wide-field imaging of the cerebellar cortex in awake, behaving mice

A major challenge in cerebellar physiology is determining how the highly conserved cytoarchitecture of the cerebellar cortex underlies its contributions to ongoing behavior and motor control. Historically, most characterizations of cerebellar neuronal activity have been limited to individual neurons or local populations within a single folium. To address this issue, we set out to develop a method for chronic, wide-field optical recordings of Purkinje cell Ca^2+^ activity in head fixed, behaving mice using transparent polymer windows, similar to those developed for the cerebral cortex^25–27^. Cerebellar windows consist of clear polyethylene terephthalate (PET) film affixed to a custom, 3D printed polymethylmethacrylate (PMMA) frame designed to conform to the natural morphology of the skull (**Figure 1A**). Combined with a head fixation implant and titanium head plate, this technique allows for chronic, wide-field cerebellar Ca^2+^ recordings in awake behaving mice (**Figure 1B**). Our cerebellar polymer windows grant optical access to a large part (∼13mm^2^) of the dorsal surface of the cerebellar cortex, including lobules 4/5, 6, and 7 of the vermis, as well as bilateral portions of the simplex and Crus I (**Figure 1C**). Importantly, this represents a substantially larger field of view than has been previously possible in chronic, awake behaving recordings, which are typically <6mm^2^. ^21–24^

**Figure 1.**
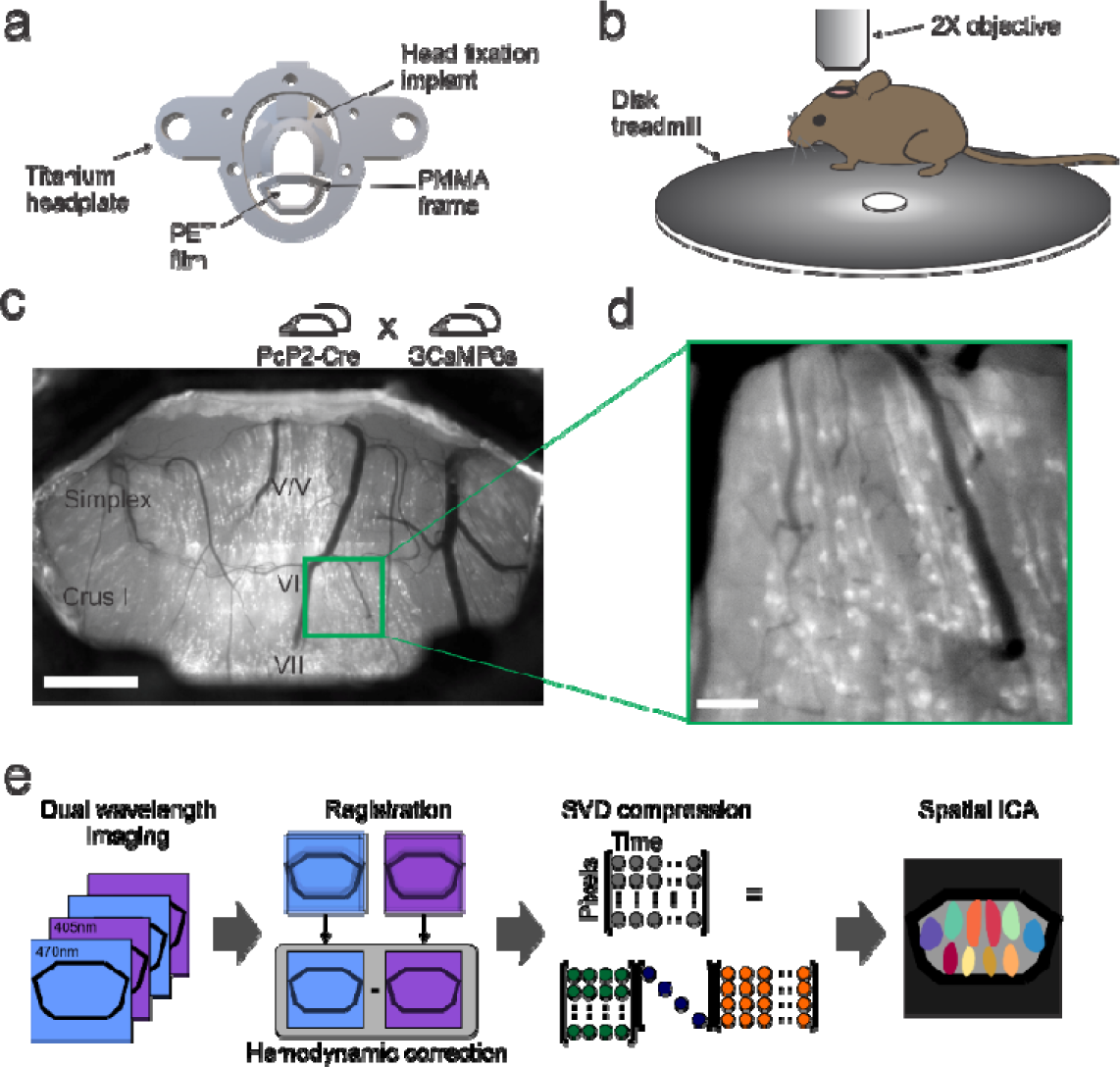
A novel method for chronic, wide-field Ca^2+^ imaging of the dorsal cerebellar cortex in awake, behaving mice. a) Cerebellar windows consist of clear PET film affixed to a custom, 3D-printed PMMA frame designed to conform to the natural morphology of the brain, and a head fixation implant attached to a titanium headplate. b) For initial imaging experiments, mice are imaged head-fixed on a freely moving disc treadmill allowing for spontaneous locomotion. c) Example wide-field field of view (FOV) in a PcP2-GCaMP6s mouse, illustrating optical access to almost the entire dorsal surface of the cerebellar cortex (scale bar 1mm). d) PcP2-GCaMP6s mice exhibit sparse expression of the Ca^2+^ indicator GCaMP6s. Combined with single-photon Ca^2+^ imaging, this allows for the resolution of individual Purkinje cell dendrites and somata (scale bar 100µm). e) Schematic of data collection and preprocessing pipeline. Dual wavelength, wide-field imaging data is collected, allowing for hemodynamic correction, which removes Ca^2+^ independent changes in GCaMP6s fluorescence. Following registration for motion correction, hemodynamic correction, and compression using singular value decomposition (SVD), functional organization of cerebellar mesoscale Ca^2+^ imaging data is achieved using the blind source separation algorithm JADER to extract distinct spatial independent components.

To visualize Purkinje cell modulation at the wide-field level, single-photon Ca^2+^ imaging was performed in PcP2-GCaMP6s mice, generated by crossing PcP2-Cre mice with floxed Ai162 mice. The resulting expression pattern in PcP2-GCaMP6s mice was sparse (**Figure 1C-D**) compared to what has been reported for PcP2-GCaMP6f crosses, with no noted alterations in Purkinje cell morphology, or widespread differences in expression patterns across cerebellar lobules, nor between animals (**Extended data 1A**). Sparse expression has been noted in other crosses with floxed lines utilizing the TIGRE locus^24^. We leveraged this sparse expression to resolve individual Purkinje cell Ca^2+^ activity at the wide-field level. Furthermore, single-photon imaging of this sparse GCaMP6s labeling allows for the recording of both Purkinje cell dendritic Ca^2+^ modulation, which is largely driven by complex spike firing, and Purkinje cell somatic modulation, which is driven more by changes in simple spike firing rate^28^ (**Figure 1D**).

### Functional segmentation of wide-field cerebellar Ca^2+^ imaging data identifies Purkinje cell dendritic and somatic networks

Wide-field Ca^2+^ imaging was performed in 8 adult mice (postnatal day 45 or greater) over a range of 3-32 weeks per animal (**Extended data 1B**) during one or more behaviors: spontaneous rest/locomotion, cued reaching, and cued reaching with prediction error. (**Figure 1E**). To achieve an unbiased segmentation of Purkinje cell network dynamics, spatial Independent Component Analysis (sICA) was used. This blind source separation analysis identifies regions of interest with maximal statistical independence and has been successfully used in wide-field Ca^2+^ imaging of the neocortex^26, 27, 29^. Importantly, this approach allows for the characterization of the functional organization of Purkinje cell activity across the cerebellar cortex irrespective of a priori assumptions about cerebellar anatomy or cytoarchitecture. Implementation of sICA at higher magnification (**Figure 2A**) reveals intriguing properties of the independent components (ICs); the ICs almost exclusively correspond to either Purkinje cell dendrites or somata (**Figure 2B-C**). Importantly, all Purkinje cell somatic and dendritic ICs were derived from a single run of sICA, with common parameters (see Methods). Dendritic and somatic ICs have clearly divergent time courses (**Figure 2B-C**, black traces), demonstrating that Purkinje cell dendritic and somatic modulation contains sufficiently distinct signals such that they are readily identified by sICA as functionally unique compartments.

**Figure 2.**
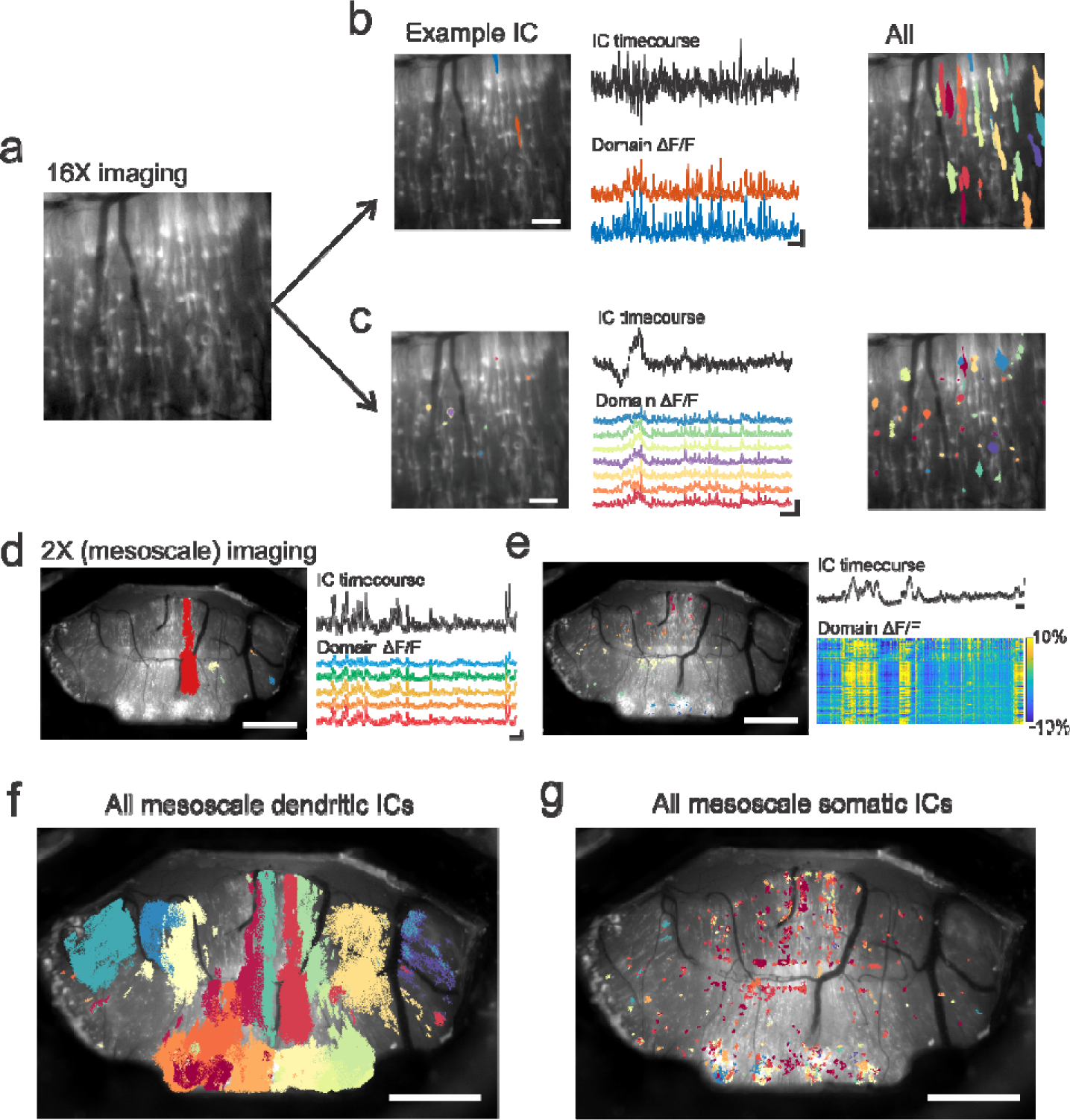
Blind source separation of Ca^2+^ imaging data segments Purkinje cell dendritic versus somatic compartments. a) Example high magnification (16X) FOV in the lobule VI of the vermis, illustrating the resolution of individual Purkinje cells with sparse GCaMP6s expression. b) Left panel: example independent component (IC) generated from sICA, which consists of two domains, both corresponding to two different Purkinje cell dendrites (scale bar: 100µm). Middle panel: Black trace: IC timecourse, red and blue traces: extracted ΔF/F from each IC domain (scale bar: 10 s, 10% ΔF/F). Right panel: all generated ICs corresponding to Purkinje cell dendrites. c) Right panel: example IC corresponding to multiple Purkinje cell somata (scale bar: 100µm). Middle panel: IC timecourse and ΔF/F for each colored somatic IC domain (scale bar: 10 s, 10% ΔF/F). Right panel: all generated ICs corresponding to Purkinje cell somata. Note the large differences in both dendritic and somatic IC timecourses as well as extracted ΔF/F from dendritic versus somatic IC domains. d) Example IC generated from wide-field FOV (2X), which consists of one large, parasagittal domain, and a smaller number of small, satellite domains. The parasagittal organization mirrors those observed previously in imaging of only Purkinje cell dendritic modulation^28^, indicating that this IC corresponds to Purkinje cell dendritic activity. Right traces are the IC timecourse and extracted ΔF/F from each IC domain (scale bar: 10 s, 10% ΔF/F). e) Example wide-field IC consisting of numerous Purkinje cell somata distributed across the imaging window. Right trace: IC timecourse for the somatic IC. Heat map is the extracted ΔF/F for each somatic domain in the IC, for the same time window shown in (d). f-g) All dendritic (f) and somatic (g) ICs at the wide-field level extracted from one example mouse during one example recording day. Each color in each panel corresponds to an individual IC; note that while dendritic ICs are largely comprised of one large primary domain, somatic ICs consist of numerous individual Purkinje cell somatic domains. Scale bars for images in D-G: 1mm.

At the mesoscale level, Purkinje cell ICs also primarily consist of two modalities with unique spatial properties. The first type of mesoscale IC consists of predominantly one large domain, often with several smaller satellite domains (**Figure 2D, F).** These ICs are organized parasagittally, and are similar to those observed during anesthetized wide-field recordings of Purkinje cells restricted to dendritic modulation^30^, suggesting these parasagittal ICs reflect dendritic modulation. During spontaneous locomotion, sICA identified 18.26 ± 2.93 (mean ± SD) dendritic ICs per recording day, with each IC consisting of 9.12 ± 6.8 domains (n = 6 mice, n = 30 recording days). The statistical independence of the different dendritic ICs suggests a functional partitioning of dendritic activity in the individual parasagittal regions. This parasagittal organization of Purkinje cell dendritic modulation is consistent with the organization of inferior olivary inputs to the cerebellar cortex^31, 32^.

Conversely, the second type of mesoscale IC consists of numerous Purkinje cell somata (i.e., domains). The domains of a given somatic IC are distributed across multiple folia, and with multiple ICs present in a folium. During spontaneous locomotion, we found 9.18 ± 3.35 somatic ICs per recording day, with 27.15 ± 21.75 domains per IC (**Figure 2E, G**). Similar to higher magnification imaging, both Purkinje cell IC modalities are yielded by the same implementation of sICA in an unsupervised fashion. This suggests that dendritic and somatic modulation of an individual Purkinje cell contains sufficiently distinct information such that the Purkinje cell belongs simultaneously to two different networks. Together, these results indicate that at the mesoscale level, Purkinje cells participate in two classes of distributed networks, with different spatial properties depending on their dendritic or somatic engagement. Dendritic Purkinje cell networks consist of large, parasagittally oriented networks, whereas Purkinje cell somatic networks consist of numerous Purkinje cell somata distributed across cerebellar folia. Given the complex and divergent spatial properties of these Purkinje cell somatic and dendritic networks identified by sICA, we set out to determine how these networks are engaged during different behaviors.

### Widespread engagement of the dorsal cerebellar cortex during spontaneous locomotion

To characterize Purkinje cell network dynamics during movement, mice were first imaged whilst head-fixed on a freely moving disk treadmill during spontaneous locomotion. Behavioral videos were labeled using DeepLabCut for markerless behavioral tracking^33^. To identify bouts of locomotion, the average broadband power (1-20Hz) X and Y position of the forepaws was computed and manually thresholded, with rapid, high magnitude increases in this average broadband power reliably identifying bouts of locomotion (**Figure 3A, Extended data Fig 3**). Spontaneous locomotion is associated with large increases in both Purkinje cell dendritic and somatic IC activity across the dorsal cerebellar cortex (**Figure 3B**), which is largely expected given the engagement of the cerebellum in motor tasks, previous observations^34–36^, as well as the various motor processes engaged during locomotion, such as limb, tail, and body movements, as well as whisking. Wide-field Ca^2+^ imaging data for all ICs was segmented using six locomotion epochs: rest, pre-walk (3 seconds prior to locomotion onset), initial walk (first 3 seconds of locomotion), continued walking, final walk (final 3 seconds of locomotion), and post walk (first 3 seconds after locomotion offset) to characterize the activity of Purkinje cell dendritic and somatic networks, as well as the interactions within these two network modalities. Widespread increases in Ca^2+^ fluorescence during the pre-walking and initial walking epochs are observed for both dendritic (**Figure 3D**) and somatic (**Figure 3G**) networks, with significant increases compared to rest beginning during the pre-walk period, and maximal increases during initial walking. Intriguingly, during continued and final walking, both increases and decreases are observed, with an overall significant increase during the post walk period (**Extended data Fig 4**).

**Figure 3.**
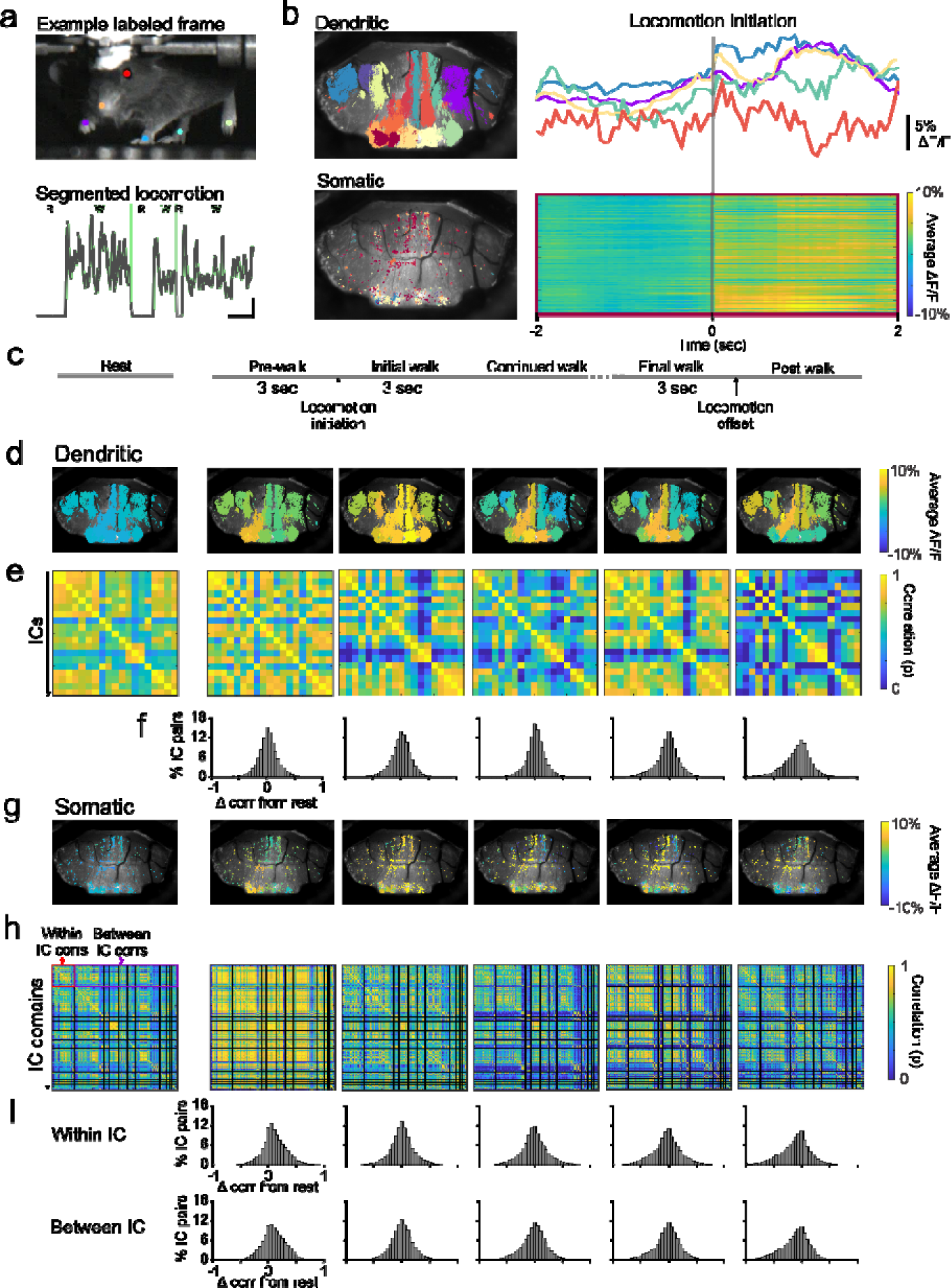
Wide-field Purkinje cell dendritic and somatic IC modulation during locomotion. a) Example frame from behavioral monitoring camera during locomotion, with body parts labeled using DeepLabCut^33^. Bottom trace: segmented locomotion times (R: rest, W: walk) identified from the average broadband (1-20 Hz) power of the extracted forepaw x and y position. This method reliably identifies periods of walking. b) Dendritic (top) and somatic (bottom) ICs from an example recording day. Right traces and heat map are the ΔF/F of example ICs (corresponding to the IC color) aligned to locomotion onset (gray bar). Somatic IC is denoted by the outline color of the heatmap. c) Imaging data is segmented for each recording day based on five locomotion epochs for subsequent analyses. d-e) Average ΔF/F and correlograms for all dendritic domains across locomotion epochs in an example animal (for one recording day). f) Population histograms of the change in correlation from rest for each IC domain across each locomotion epoch (n = 6 mice, n = 30 recording days). g-i) Same as d-f but for somatic networks. Note that since somatic ICs consist of numerous domains, both within IC and between IC correlation changes are quantified across all domains, in which within IC changes denote comparisons across somatic domains belonging to the same IC, whereas between IC changes denote comparisons across somatic domains belonging to different ICs.

**Figure 4.**
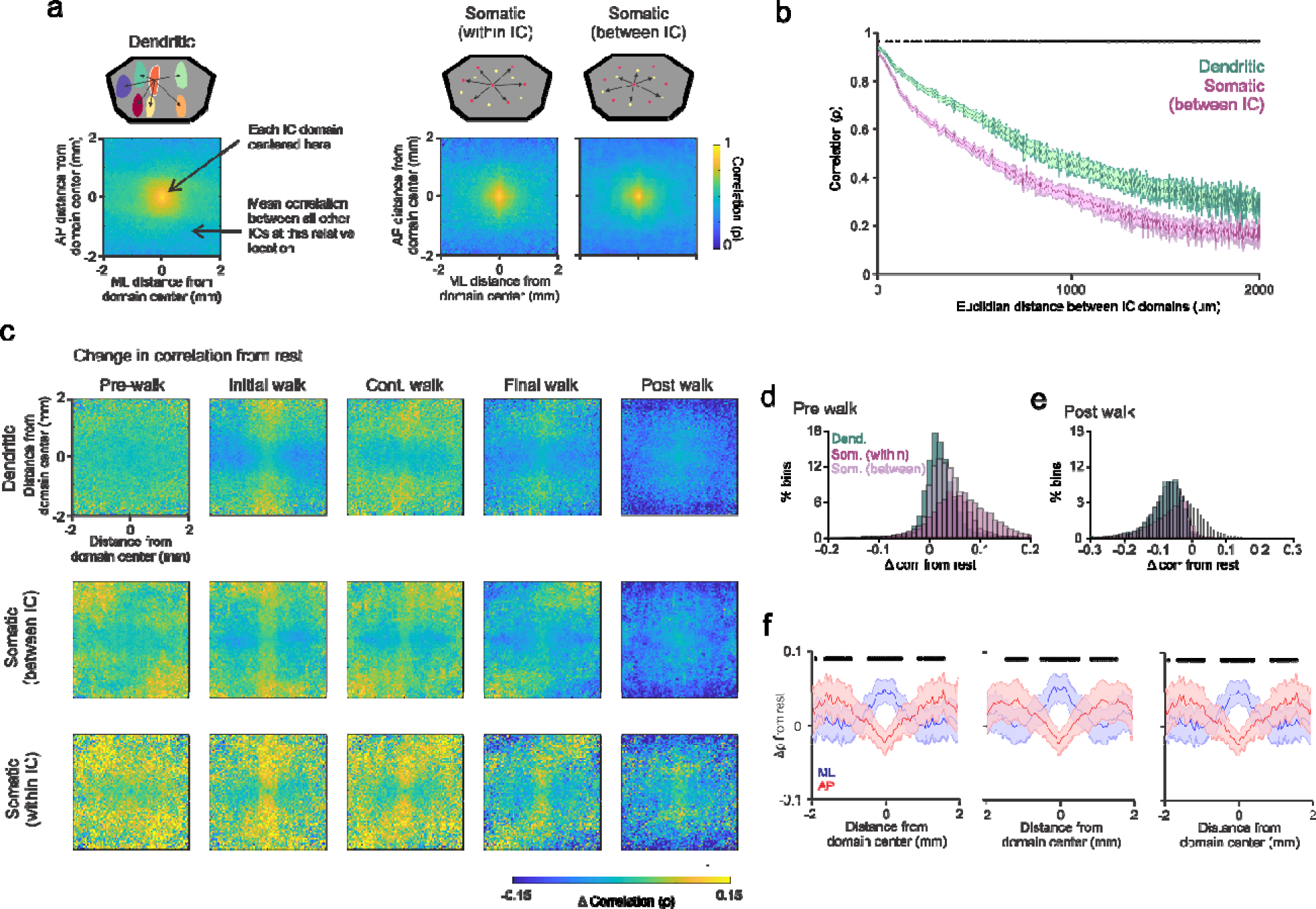
Changes in spatial correlations for Purkinje cell dendritic and somatic ICs during locomotion epochs. a) Spatial correlation analysis schematics (top) and spatial correlation maps (bottom) at rest for dendritic versus somatic (within IC and between IC) networks. b) Average (mean ± SEM) correlation (ρ) across all mice, all recording days, as a function of distance between IC domains for dendritic ICs (green trace) versus somatic ICs (purple trace, between IC domain correlations). Somatic ICs show a significant decrease as a function of distance compared to dendritic ICs (black dots over traces denote p < 0.05 t-test with false discovery rate correction [Benjamini & Hochberg (1995)]). c) Population change in spatial correlation maps compared to rest for dendritic IC domains, somatic within IC domains and between IC domains across all locomotion epochs. d) Histogram of population change in spatial correlation map bins for the pre-walk period. Somatic IC domains demonstrate significantly increased correlations throughout the spatial map during pre-walk compared to dendritic domains (One-way ANOVA F(2,19192) = 1581, p < 0.001. Tukey HSD, p < 0.05 for dendritic IC versus somatic within and between IC). e) Same as for (d), but for post walk periods. Dendritic IC domains demonstrate significantly decreased correlations throughout the spatial map during post walk periods compared to both somatic within and between IC correlations (One-way ANOVA F(2,19192) = 741.61, p < 0.001. Tukey HSD, p < 0.05 for dendritic IC versus somatic within and between IC. f) Average (mean ± SEM) change in correlations compared to rest for initial walking along the mediolateral (blue) and anteroposterior (red) axes across all mice illustrate alternating parasagittal bands of increases and decreases in correlations emerging during locomotion (black dots: p < 0.05 t-test with false discovery rate correction^54^.

As we were not only interested in changes in Ca^2+^ fluorescence, but also the patterns of correlations in the Purkinje cell dendritic and somatic networks, we measured the changes in correlation of Ca^2+^ fluorescence across ICs during locomotion epochs. For dendritic ICs, the correlation between each IC was computed. For somatic ICs, as these ICs consist of numerous domains, we measured the correlations for domains belonging to the same IC (within IC correlations) as well as the correlations for domains belonging to different ICs (between IC correlations). Intriguingly, the patterning and degree of dendritic and somatic IC domain correlations change over the different periods and often diverge from the changes in overall fluorescence. For dendritic networks, modest increases in the average correlations begin during pre-walking periods (change in correlation (ρ) from rest: 0.017 ± 0.18 (SD), p < 0.001, student’s t-test) and there is a decrease in overall correlations during the post-walk period (change in correlation from rest: - 0.07 ± 0.24 (SD), p < 0.001, student’s t-test) (**Figure 3E-F**). More importantly, the correlations undergo a non-uniform redistribution among domains (**Extended data figure 5A)**. For both somatic within and between IC correlations, locomotion preparation (the pre-walk period) evokes broad increases in somatic correlations (change in correlation from rest: 0.11 ± 0.21 within IC; 0.096 ± 0.21 between IC, for each p < 0.001, student’s t-tests). Similar to dendritic ICs, the pattern of correlations for somatic networks changes throughout the course of locomotion epochs (**Extended data figure 5B**), concluding with a significant decrease at locomotion offset (change in correlation from rest: -0.12 ± 0.27 within IC; -0.12 ± 0.26 between IC, p < 0.001, student’s t-tests) (**Figure 3H-I**).

**Figure 5.**
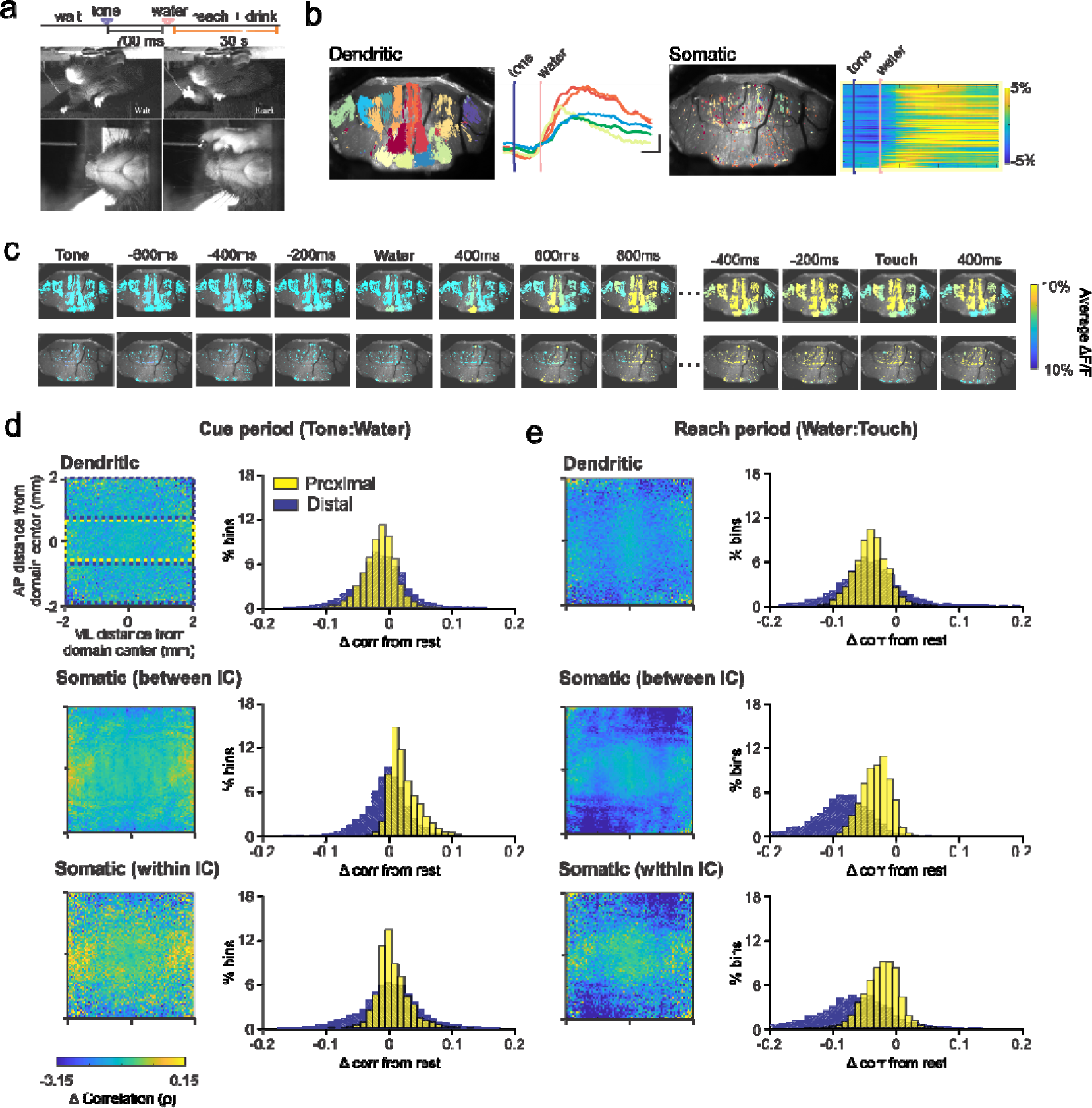
Widespread activation of Purkinje cell somatic and dendritic networks, with dynamic transverse alterations in somatic network correlations during cued reaching. a) Schematic of cued reaching paradigm. Mice are trained to make targeted reaches towards a water spout. In each trial, an auditory tone cues the delivery of a water droplet from the spout. The mouse subsequently reaches for the reward. Reach trials are separated by 30 seconds. b) Dendritic (left) and somatic (right) ICs and example averaged ΔF/F across 40 reach trials. Blue bar: timing of auditory cue, pink bar: water droplet. For somatic IC, ΔF/F for each domain is displayed as a heat map as in Fig 2B. c) Averaged ΔF/F maps for all dendritic (top) and somatic (bottom) ICs aligned to auditory cue, water droplet delivery, and times in which the mouse made contact with the water spout (touch). d) Left panels, change in spatial correlation maps compared to rest for the auditory cue period for dendritic and somatic (within and between) IC domain correlations (n = 4 mice, over recording days). Right histograms denote the change in correlation for population map bins in the proximal third (yellow) versus distal 1/3 (blue) of each map in the anterior-posterior dimension. No significant difference in spatial correlations is observed for proximal versus distal dendritic IC domains (proximal change in correlation: -0.013 ± 0.024 vs distal change in correlation: -0.015 ± 0.04; p = 0.08, student’s t-test), whereas significantly increased correlations are observed for proximal somatic IC domains compared to distal (between IC proximal change in correlation: 0.024 ± 0.022 vs distal change in correlation -0.0032 ± 0.032; within IC proximal change in correlation: 0.017 ± 0.025 vs distal change in correlation: 0.0013 ± 0.046, p < 0.001, student’s t-tests). e) Same as for (d), but for the reaching period. Whilst all differences in proximal versus distal correlations are statistically significant (p < 0.001, student’s t-test), stronger decreases in distal domains are observed for somatic IC domains (between IC proximal change in correlation: -0.031 ± 0.022 vs distal change in correlation: -0.092 ± 0.055; within IC proximal change in correlation: -0.023 ± 0.026 vs distal change in correlation: -0.071 ± 0.067, p < 0.001, student’s t-test).

Together, these results indicate that while large regions of the cerebellar cortex are highly activated during behavior, these overall activity levels can be associated with highly differing correlation dynamics across Purkinje cell networks. Next, we sought to determine if and how spatial arrangements of Purkinje cell networks impact correlation dynamics. To do this, we computed spatial correlation maps for all dendritic and somatic IC domains (see Methods). These spatial correlation maps characterize the average correlation between IC domains as a function of their relative spatial positioning for dendritic IC domains and somatic IC domains (both within IC and between IC) (**Figure 4A**). At rest, both dendritic and somatic ICs have high spatial correlations for more proximally located domains, which decrease as the distance between IC domains increases. However, somatic IC domains demonstrate slightly more restricted spatial correlations, as the correlation between IC domains significantly decreases with distance compared to those for dendritic ICs (**Figure 4B**, black dots indicate p < 0.05 with false discovery rate correction).

We next quantified the changes in spatial correlations during the different locomotion epochs compared to rest and found behavior-dependent alterations that primarily reflect the two major architectures of the cerebellar cortex, parasagittal banding of the olivocerebellar projection and the transverse projections of the parallel fibers (**Figure 4C**). In agreement with the broad IC correlation analyses, the pre-walk period is associated with significantly increased overall correlations for somatic and dendritic domains, but is the strongest for somatic networks (**Figure 4D**, One-way ANOVA F(2,19192) = 1581, p < 0.001. Tukey HSD, p < 0.05 for dendritic IC versus somatic within and between IC), and the post-walk period is associated with broad decreases in correlations for both dendritic and somatic ICs. However, during locomotion, the spatial correlation analysis reveals that the increases and decreases in correlations observed contain a particular spatial organization, such that alternating parasagittal bands of correlation changes emerge within this 4mm^2^ spatial region (**Figure 4F**, black dots indicate p < 0.05 with false discovery rate correction). For example, during the initial walk, both the dendritic and somatic ICs show an anterior-posterior (e.g. parasagittal) band of increased correlation compared to rest and decreases in correlations in the medial-lateral (e.g. transverse) direction (**Figure 4C**). The magnitude and timing of these increases depends on type of IC, with the dendritic and between somatic ICs showing the strongest transverse decrease, whereas for the within somatic ICs, the increases in the parasagittal direction are most prominent. The emergence of these parasagittal and transverse bands during locomotion indicates that this behavior is associated with changes in Purkinje cell network spatial correlations, which largely follow the established architecture of the cerebellar cortex.

### Characterizing cerebellar modulation during a cued reaching task

We next sought to determine how Purkinje cell somatic and dendritic network dynamics are altered during learned behaviors. Therefore, wide-field Purkinje cell Ca^2+^ imaging was performed in mice trained in a cued reaching task, in which an auditory tone cues the delivery of a water droplet at the end of a spout. Mice subsequently reach for the water droplet and bring it to their mouths to drink (**Figure 5A**). Well-trained mice perform the task at a high level of consistency and skill, exhibiting 90% reaching accuracy over greater than 50% of recording days (**Extended data figure 6A**), and the majority of reaches taking place within 2 seconds after water droplet delivery (**Extended data figure 6B**). As with locomotion, widespread, bilateral activation of both Purkinje cell dendritic and somatic ICs occurs during reaching (**Figure 5B-C**). The activation emerges shortly after the delivery of the water droplet and continues throughout the duration of the reach itself.

**Figure 6.**
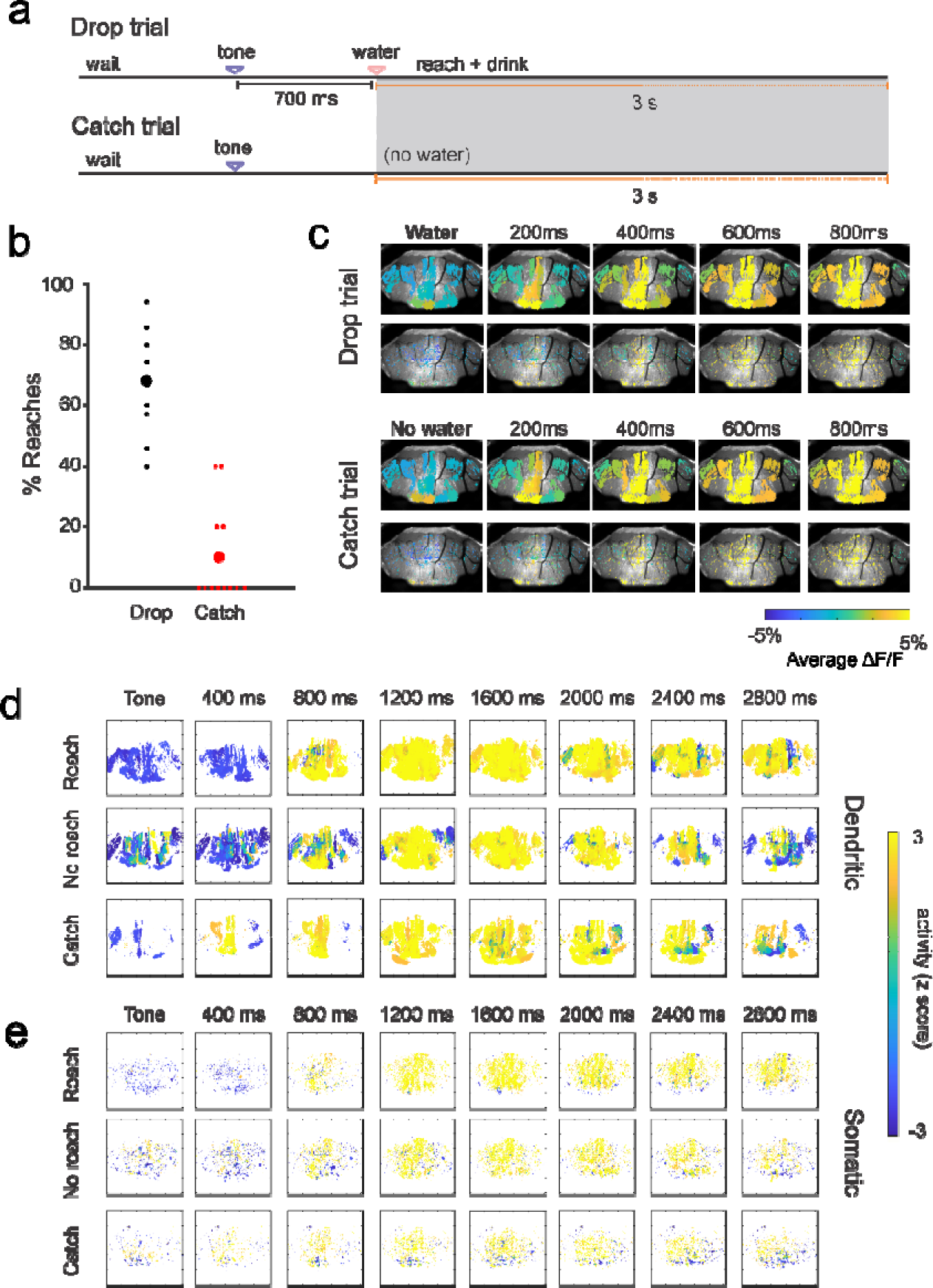
Widespread activation of the cerebellar cortex during different reaching paradigms and behavioral outcomes. a) Schematic of prediction error paradigm. Reach trials either consist of standard cued reaches, with auditory tone followed by water droplet delivery, or catch trials, in which the water droplets omitted (catch trials). Catch trials are administered randomly at a frequency of once every eight reach trials (12.5% probability). b) Probability to reach for the spout for water droplet (black) versus catch (red) trials across all mice and recording days (n = 4 mice, over 12 recording days). In > 75% of recording days, mice did not make a single reach during the catch trial. c) Averaged ΔF/F maps for all dendritic and somatic ICs aligned to the timing of expected water droplet delivery in an example animal. Note that widespread cerebellar engagement occurs during both conditions. d-e) Population maps (n = 4 mice, over 30 total recording days) of thresholded, normalized modulation of dendritic and somatic networks aligned to the auditory cue (0 ms) during trials in the standard reaching paradigm in which the mouse successfully reached for the spout (“Reach”), trials in which the mouse failed to reach for the spout (“No reach”), and during the catch trial paradigm in which the water droplet was omitted (“Catch”) illustrate a high degree of cerebellar activation in all three conditions.

Spatial correlation analyses reveal divergent effects of both the auditory cue as well as the reach on dendritic versus somatic networks. During the cue period (the time at which the auditory cue is delivered up until water droplet delivery), the spatial correlation maps for the dendritic networks change little, including the spatial pattern, compared to the rest period (**Figure 5D**, top map and histogram). Somatic networks, however, show increased correlations, which are strongest along the transverse zone (**Figure 5D** middle and bottom maps and histograms). During the reach period (the time from water droplet delivery to the time the mouse touches the spout for reaches within 2 s after droplet delivery), broad decreases in correlations are observed for both somatic and dendritic ICs (**Figure 5E**). However, similar to the cue period, the magnitude of these decreases depends on relative anterior/posterior positioning for somatic domains, with the strongest decreases in distal domains (e.g. off of the approximate transverse zone). Together, these results demonstrate that while spontaneous locomotion and reaching evoke similar, widespread activation of the cerebellar cortex, the underlying spatial correlation patterns are markedly different. For somatic networks, Purkinje cells more proximal in the anterior/posterior dimension (i.e. within the theoretical transverse zone) show increased correlation in Ca^2+^ modulation during the presentation of an auditory cue (**Figure 5D**, bottom two rows) followed by subsequent decreases in correlations for the distal domains (i.e. between transverse zones) during the reach itself (**Figure 5D**, bottom two rows). These changes in correlations contrast with those observed for spontaneous locomotion, demonstrating that different motor behaviors engage and modify Purkinje cell correlation dynamics in distinct ways, though in both cases, the modifications obey overarching structural organization(s) of the cerebellar cortex.

### Prediction errors trigger widespread changes in somatic networks

As error processing and prediction are considered hallmarks of cerebellar function, we next examined how prediction errors impacted Purkinje cell population dynamics. To do this, catch trials were implemented during the reaching paradigm, in which one out of every eight auditory cues in a given imaging stack was randomly selected to not result in water delivery (**Figure 6A**). Mice seldom executed a reach during the catch trial, with >75% of recording days yielding zero reaches during catch trials (**Figure 6B**). In addition to examining prediction error processing, this also allowed for a determination of how Purkinje cell activity and correlation dynamics are altered in the absence of a reach. Both Purkinje cell dendritic and somatic ICs are activated during water drop trials, when the animal reached for the water droplet, and catch trials, when no water was delivered, and the animal did not reach (**Figure 6C**). Across the population, widespread activation of the cerebellar cortex occurs for vastly different reaching outcomes: during the standard reaching paradigm, when the mouse successfully reached (dendritic ICs, peak change in activation (z-score) 4.19 ± 1.97; somatic ICs, peak change in activation 3.99 ± 1.48), when the mouse failed to reach (dendritic ICs: 4.13 ± 1.78; somatic ICs: 4.13 ± 1.58), and during the catch trial when no water was delivered (dendritic ICs: 3.61 ± 1.18; somatic ICs: 3.76 ± 1.16) (**Figure 6D-E**, n = 4 mice). In all three cases, widespread, bilateral activation emerges shortly after the auditory cue, and persists throughout the duration of the potential reaching period, whether the animals receive water and simply fail to reach, or whether there is no water present. This is true for both the dendritic (**Figure 6D**) and somatic networks (**Figure 6E**), however there are some differences, such as less peak activation during the catch trial, particularly for dendritic networks (One-way ANOVA F(2,2991) = 23.11, p < 0.001. Tukey HSD, p < 0.05 for both reach and no reach versus catch peak change in activation). Overall, the pattern of activation at this wide-field level is not a particularly informative representation of the condition or the reach itself (as increases happen equally in reach and no reach trials).

Given the similarities in activation during the reach, no reach and catch trial conditions, we next investigated whether the spatial correlations were altered during the catch trial. Spatial correlation analyses reveal extensive changes in correlations during the catch trial, particularly for somatic networks. Broad increases in correlations for Purkinje cell somatic networks are evoked during the expected water delivery period in catch trials compared to water drop trials (**Figure 7A**). These catch trial associated increases in somatic IC correlations, both within and between IC domains, are significantly greater than those in dendritic networks (**Figure 7B**, One-way ANOVA, F(2, 19113) = 5299, p < 0.001. Tukey HSD, p < 0.05 for somatic within, between IC vs. dendritic between IC change in correlation). While increased correlations are observed across virtually the entire somatic IC spatial correlation maps, they appear strongest off the transverse zone. Therefore, prediction errors trigger increases in Purkinje cell somatic correlation that are influenced in part by the transverse organization, but spatially very widespread, highlighting that errors have the capacity to dramatically influence somatic network dynamics.

**Figure 7.**
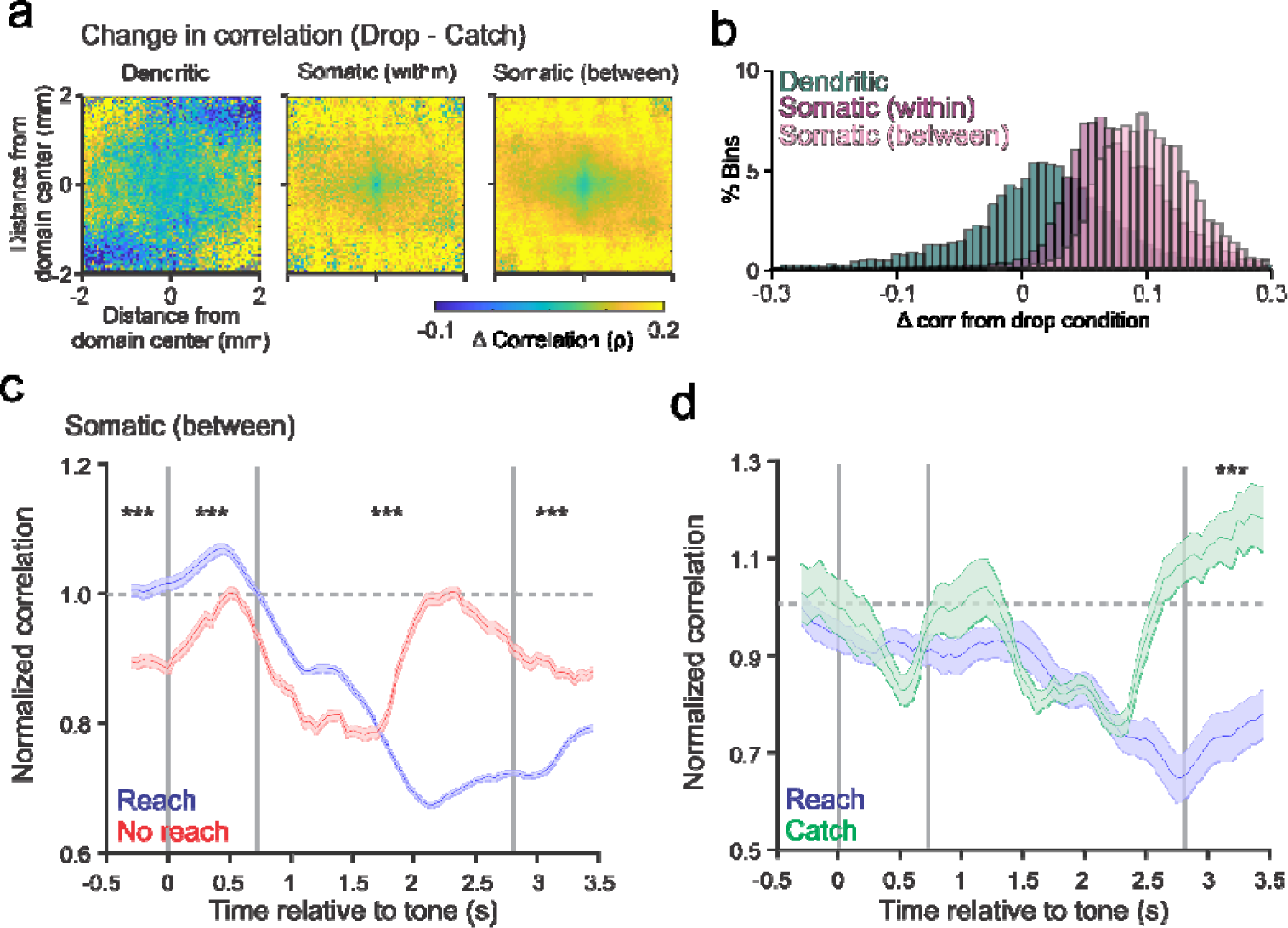
Behavioral outcome dependent alterations in somatic network interactions during the catch trial. a-b) Population changes in spatial correlations during the catch trials relative to the drop condition illustrate dramatic increases in correlations for somatic networks compared to dendritic networks (n = 4 mice, over 12 recording days). b) Histograms of change in spatial correlations for dendritic and somatic (both within IC and between IC) domains illustrate significantly increased correlations for somatic IC domains compared to dendritic domains (One-way ANOVA, F(2, 19113) = 5299, p < 0.001, Tukey HSD, p < 0.05 for both somatic within and between IC vs. dendritic between change in correlation). c) Average normalized correlations (relative to baseline, see Methods) between somatic IC domains (between IC) relative to auditory cue for reach (blue) and no reach (red) conditions during the standard reaching paradigm. Eventual successful reaches are associated with increased correlations during the pre-cue periods compared to those for failed reaches (One-way ANOVA, F(1,13137) = 107.47, p < 0.001). Successful reaches are also associated with significantly higher somatic IC domain correlations during the cue period compared to failed reaches (One-way ANOVA, F(1,13137) = 55.9, p < 0.001). During the reach period, whilst both reach and no reach outcomes are associated with initial decreases in correlations, failed reaches demonstrate a return to baseline correlation whereas successful reaches show a further decrease that is maximal prior to the thresholded maximum reach time (2 seconds after water droplet delivery), resulting in a significant overall decreased compared to failed reaches (One-way ANOVA, F(1,13137) = 293.17, p < 0.001). d) During the catch trial paradigm, somatic network correlations are significantly increased after the time of anticipated water delivery compared to trials in which water was delivered and mice reached successfully (One-way ANOVA, F(1,9604) = 28.66, p < 0.001).

Given the large changes in correlation during the different reaching epochs in Purkinje cell somatic networks, we sought to determine whether the time course of these patterns depended on whether the mouse executed a successful reach versus did not reach. To do this, we quantified the change in correlation compared to rest across all somatic ICs over time using a sliding window analysis in our standard reaching paradigm. For trials in which mice failed to reach despite the auditory tone and water droplet delivery, somatic IC domain correlations are significantly decreased just prior to the onset of the auditory cue (**Figure 7C**, One-way ANOVA, F(1,13137) = 107.47, p < 0.001), suggesting that hypo-correlated activity of somatic networks is associated with the failure to ultimately execute the movement. In line with this interpretation, during the cue period, the correlation between somatic IC domains is significantly increased in trials in which the mouse subsequently successfully executed a reach compared to failed reach trials (One-way ANOVA, F(1,13137) = 55.9, p < 0.001). During the reach period, an intriguing time course is observed. Somatic IC correlations decrease initially in both conditions. However, there is an inflection point at which the correlation magnitude in failed reach trials returns to baseline, whereas for the reach trials the correlation continues to decrease during the reach period, resulting in significantly decreased somatic IC correlations for reach versus no reach outcomes (One-way ANOVA, F(1,13137) = 293.17, p < 0.001). Therefore, somatic networks not only modulate their correlation dynamics during a movement, but also their dynamics prior to movement may contribute to the likelihood of ultimately successfully executing that movement.

In contrast to the increase in somatic correlations seen for successful reach trials in the standard reaching paradigm, during the catch trial paradigm, trials in which the water was delivered and mice subsequently successfully execute a reach are not associated with an increase in somatic IC correlations during the auditory cue period (**Figure 7D**), perhaps reflecting the lack of certainty in the association between the auditory cue and water delivery. The somatic IC correlations are similar for the reach and catch trials until late in the reach period, at which point there is a significant increase in correlation during catch trials (One-way ANOVA, F(1,9604) = 28.66, p < 0.001). Similar increases in correlations occur for somatic within IC changes in correlations (**Extended data Fig 7**). Dendritic ICs correlations also significantly decrease during reach versus no reach trials, though changes in the correlation during the auditory cue period fail to predict reaching outcome (**Extended data Fig 7**). During the catch trial paradigm, the impact of task outcome on dendritic correlations is less clear than for the somatic networks, with differences observed prior to auditory cue and the timing of water droplet delivery (**Extended data Fig 7**). Therefore, while the correlations for dendritic ICs modulate with some of the different behaviors, the dendritic network dynamics are not likely driving the somatic network correlation changes. Together, these results demonstrate that somatic networks alter their interactions independent of the levels of activation, with the magnitude of somatic network correlations prior to movement linked to the eventual behavioral outcome, and prediction errors evoking large increases in correlations.

## Discussion

Despite decades of research, a clear link between cerebellar architecture and function, especially with respect to Purkinje cell dynamics, has proved elusive. Our results demonstrate that Purkinje cells simultaneously belong to multiple, distinct networks - one based on their somatic modulation and one their dendritic modulation, which are likely driven by their simple spike versus complex spike activities, respectively. These networks have unique spatial properties, with dendritic modulation corresponding to more spatially clustered, parasagittal bands, whereas somatic modulation consists of functional populations of Purkinje cells across folia throughout the dorsal surface of the cerebellar cortex. These somatic and dendritic networks also differ in their patterns of activation and correlation, with changes in the correlation dynamics often following the overarching structural organization of the cerebellar cortex. For the tasks evaluated, the spatial correlation dynamics are the strongest for somatic networks during cued reaching, with changes in somatic network correlations (but not their overall activity levels) reflective of ultimate behavioral outcomes. In this view, Purkinje cell functional networks can be parsed by their activity modalities (i.e., dendritic versus somatic modulation), the spatial correlation profiles of those networks are dynamic during behavior, with changes following the overarching structural organization of the cerebellar cortex and reflective of behavioral states and task demands.

Most previous optical imaging approaches have been limited to recording local populations of neurons within either a folium or < 6mm^21–24^, and have utilized optical imaging techniques and expression strategies restricted to recording Purkinje cell dendritic modulation. While a recent study performed wide-field Ca^2+^ imaging of the cerebellar cortex over a similar spatial scale as described here, the imaging was also restricted to dendritic modulation and was largely performed in anesthetized mice^30^. To our knowledge, the present study provides the largest field of view of Purkinje cell modulation in awake, behaving animals. This is in part due to the design of our cerebellar windows that conform to the natural contours of the skull, minimizing any deformation that can occur with e.g. glass coverslips. In addition, our single-photon imaging approach with sparse GCaMP6s expression provides for a novel characterization of Purkinje cell network dynamics with respect to both somatic and dendritic modulation, simultaneously.

We did not anticipate that sICA would result in such a marked division of Purkinje cell Ca^2+^ fluorescence into two classes of ICs, composed almost exclusively of either Purkinje dendrites or somata. As sICA is an unbiased, blind source separation method for identifying statistically independent regions from a complex mixture of sources and has no information or assumptions about the underlying cellular or circuit architecture, this suggests the division of Purkinje cells into functional dendritic and somatic networks is a fundamental organizing principle. The nearly complete separation of dendritic versus somatic ICs, their presence in all animals and behaviors across recording days, and the differences in the modulation and correlation patterns at different behavioral epochs further support this concept. While the parasagittal dendritic ICs, which have been observed previously with electrophysiological and optical recordings, undoubtedly reflect the well-known olivo-cerebellar projection of the climbing fibers^8^ and their action on Purkinje cells, we can only speculate on the multiple folia distribution of the somatic ICs. Several factors are likely to contribute to the location of the somata across multiple folia, including the transverse organization of parallel fiber inputs, the widely distributed mossy fiber input into the cerebellar cortex^17^, and synaptic plasticity^14, 15^.

One common hypothesis asserts that cerebellar neurons are characterized by sparse coding of behavior^37–40^. The present findings do not support a sparse coding viewpoint, as there is widespread bilateral activation of Purkinje cells during the three behaviors: locomotion, reach, and catch trials. While not unexpected for locomotion that engages all four limbs as well as much of the body, widespread activation was not necessarily anticipated for reaching that is restricted to a forelimb. Finding widespread activation in the cerebellum in the reach task, even in trials where the animal does not reach, was particularly surprising. While unexpected, this widespread activation largely mirrors recent cerebral cortical observations, using either wide-field imaging or large scale, multiple electrophysiological recordings, which describe widely distributed encoding of behavior across brain structures and circuits^29, 41–45^. As different cerebellar regions have differing downstream connectivities, this widespread engagement of the cerebellum also suggests widespread cerebellar influence on those downstream areas during these behaviors. The behavior-related changes in spatial correlation structure among Purkinje cells also suggests a degree of functional coordination across these areas, which is dynamic and depends on behavioral state and task demands but has a spatial pattern reflective of the parasagittal and transverse organization of the cerebellar cortex.

A common feature of the network level Purkinje cell dynamics characterized here is the seemingly divergent impacts of behavior on the levels of network activity versus correlated activity within the network. During the reaching paradigms, overall network activity levels differs remarkably little between trials with a successful reach versus trials where no reach was executed. In contrast, the spatial patterns of correlation in somatic networks are dynamic across periods within a reach trial, and correlation patterns differ across trials with a successful reach versus trials without a reach. The correlation profiles even differ between successful reach trials during the standard reaching task and successful reach trials during the catch trial paradigm, despite similar behavioral outcomes, suggesting state dependency (e.g. the predictive value of a tone). In addition, the hypo-correlation of somatic networks prior to the auditory cue when the mouse ultimately does not reach in the standard paradigm suggests that the state of cerebellar network dynamics is important for execution of cued behaviors and potentially reflects one or more factors, including attention, motivation, or expectations. In short, the spatial patterns of correlation in somatic networks is highly linked to information about task cues, the execution of a reach, and prediction error. Furthermore, these correlations can be linked to the ultimate behavioral outcomes, such as the failure to execute a reach.

While at the spike level, the potential functional role(s) of synchronous Purkinje cell output is a topic of continued debate^46–52^, the correlated modulation of somatic activity described here likely reflects overall simple spike modulation rather than conveying information about spike level dynamics, due in part to the kinetics of GCaMP6s^53^ and the high firing rates of Purkinje cells^28^. Given the convergence of Purkinje cells onto neurons in the deep cerebellar nuclei, which ultimately project to numerous downstream regions, one possibility is that these different correlation patterns induce distinct output patterns of DCN neurons. In this view, the spatial complexity of somatic networks, and their capacity for intricate spatial correlation patterns might allow for information rich interactions between Purkinje cell modulation and their ultimate influence on DCN output.

Using wide-field imaging, the results reveal a strong connection between cerebellar cortical structure and function, in which the spatially distinct Purkinje cell dendritic and somatic networks uncovered dynamically change their correlation patterns. These correlation pattern changes are reflective not only of behavior, but also potentially internal state, and display spatial properties likely supported by the cytoarchitecture of the cerebellum. Importantly, uncovering of these correlation dynamics among Purkinje cells widely distributed across folia in awake behaving animals required our innovative imaging approach. Together, our observations highlight that the dynamic correlations across Purkinje cell networks, rather than just overall activity levels, is a key feature of the cerebellum’s contributions to behavior.

## Acknowledgments

This work was supported in part by NIH K99NS121274 (MLS), P30 DA048742 (TJE and SBK), an American Epilepsy Society Postdoctoral Fellowship (MLS), NIH R01-NS112518 (EKM), NIH R01-NS111028 (TJE and SBK), and a University of Minnesota McKnight Land-Grant Professorship award (EKM).

## Author contributions

MLS, REC, and TJE designed the behavioral paradigms. SBK, VR, MLS, REC, and TJE conceived of and designed the implant design. MLS, BWK, KT, and EW collected the data. MLS analyzed the data, with input from TJE and EKM. MLS and TJE wrote the manuscript.

## Extended data figures

**Extended data figure 1.**
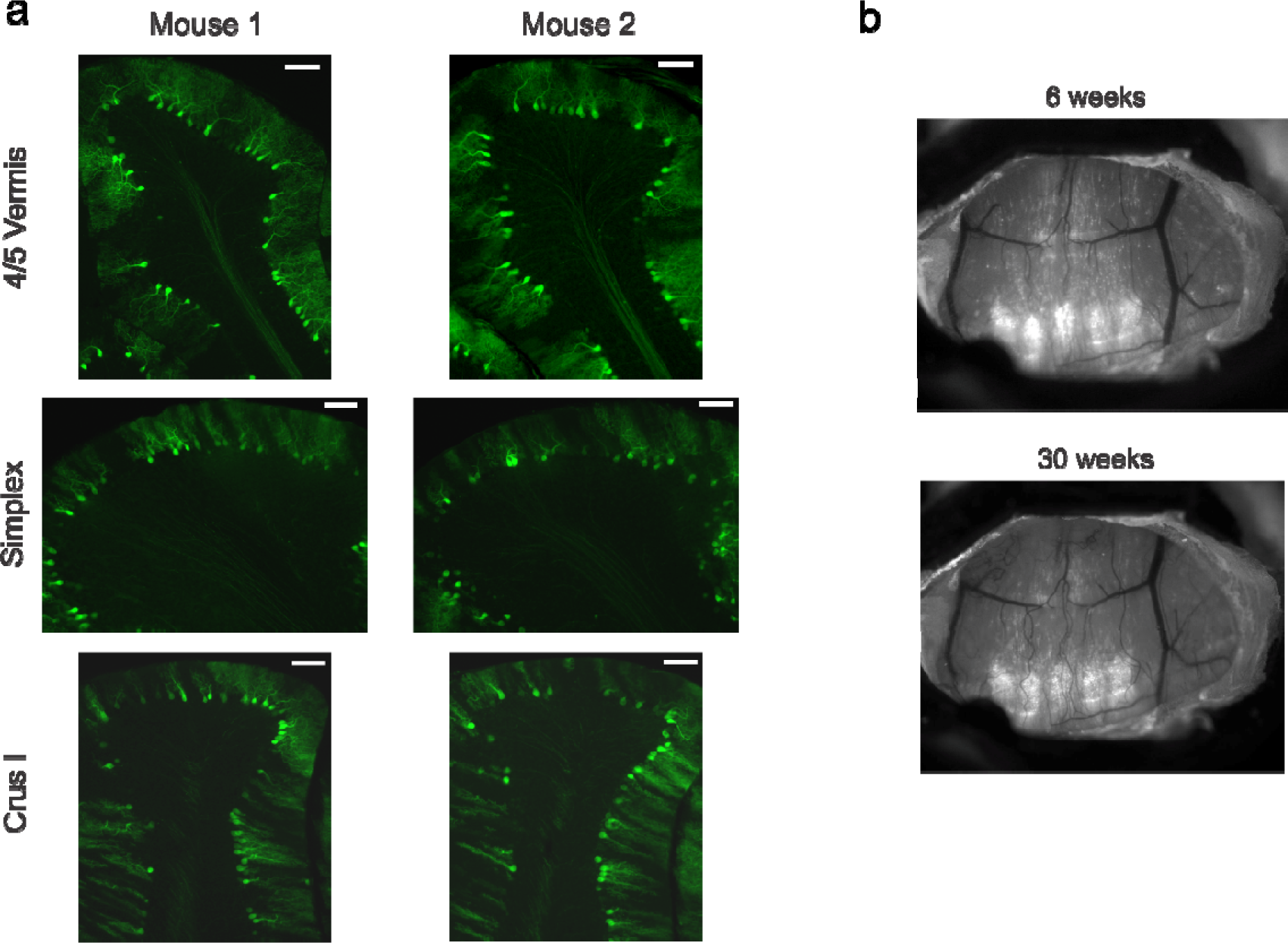
Chronic wide-field imaging of sparsely expressed GCaMP6s in Purkinje cells. a) Sagittal slices from two example mice illustrating sparse GCaMP6s expression throughout the lobules within the imaging field of the cerebellar window. Scale bars: 100µm. b) Imaging field of an example mouse at 6 weeks post implant (top) and 30 weeks post implant (bottom) illustrating that cerebellar windows are effective for chronic imaging applications.

**Extended data figure 2.**
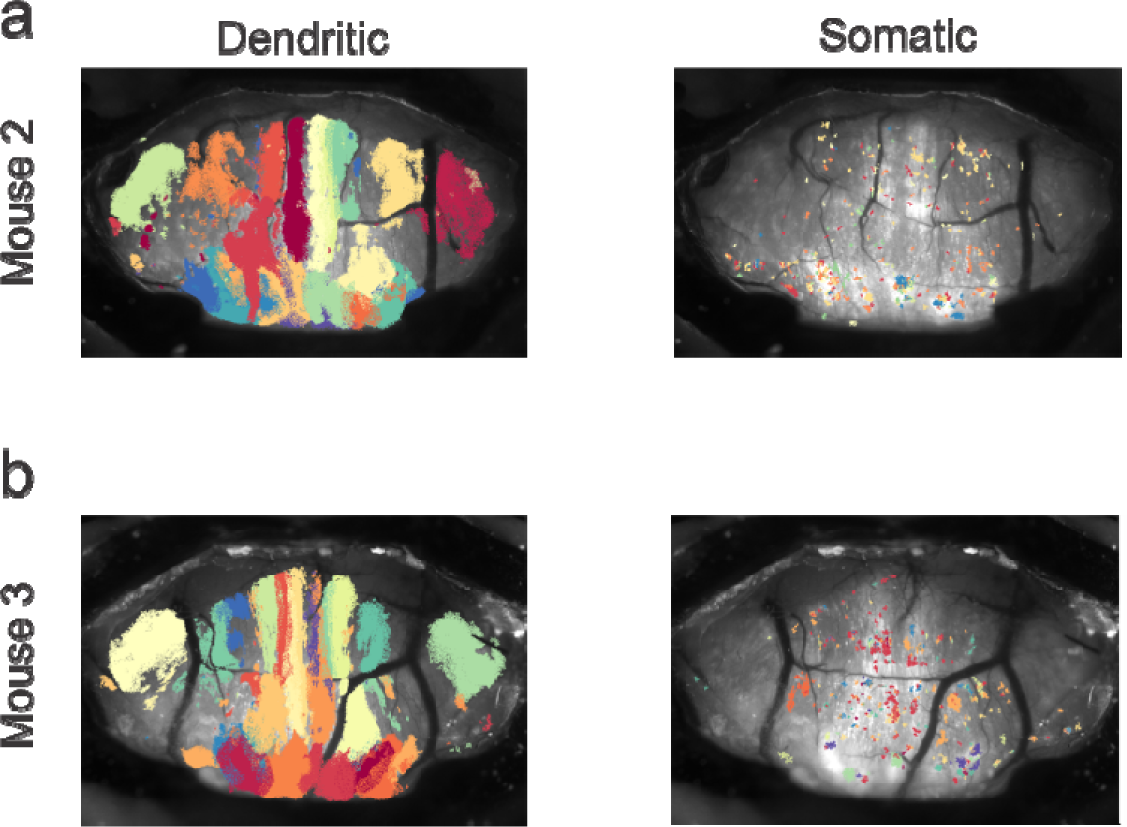
Dendritic and somatic ICs across mice. a-b) Example dendritic (left) and somatic (right) ICs in two additional example mice.

**Extended data figure 3.**
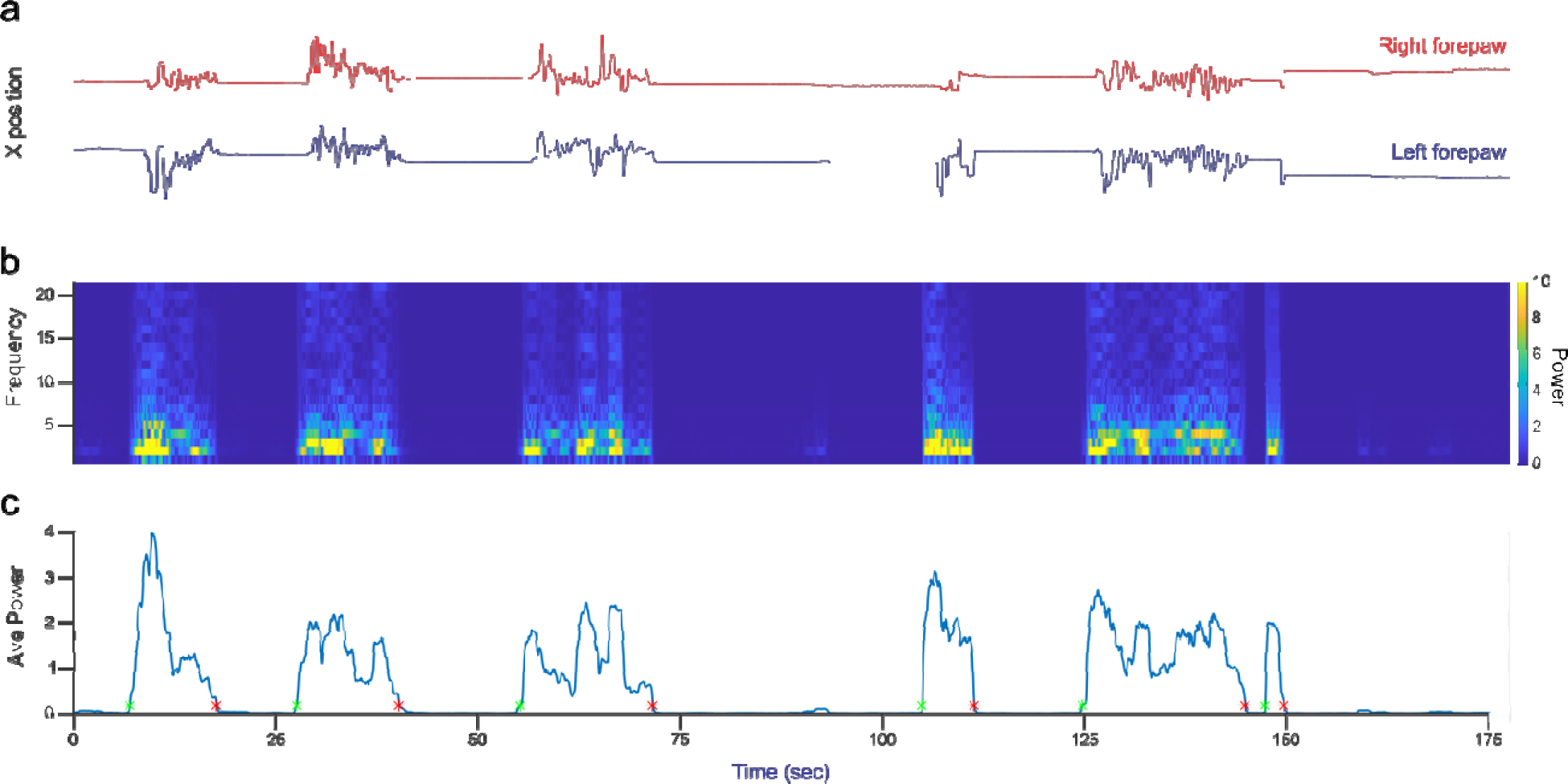
Segmentation of locomotion periods from forepaw position. a) Extracted X position from the right (red) and left (blue) forepaw using DeepLabCut from a mouse on the disk treadmill. b) Spectrogram of frequency of X position over time, illustrating increases in broadband power during locomotion. c) Average power in the 1-20 Hz range over time, with thresholded periods of locomotion onset (green dots) and offset (red dots).

**Extended data figure 4.**
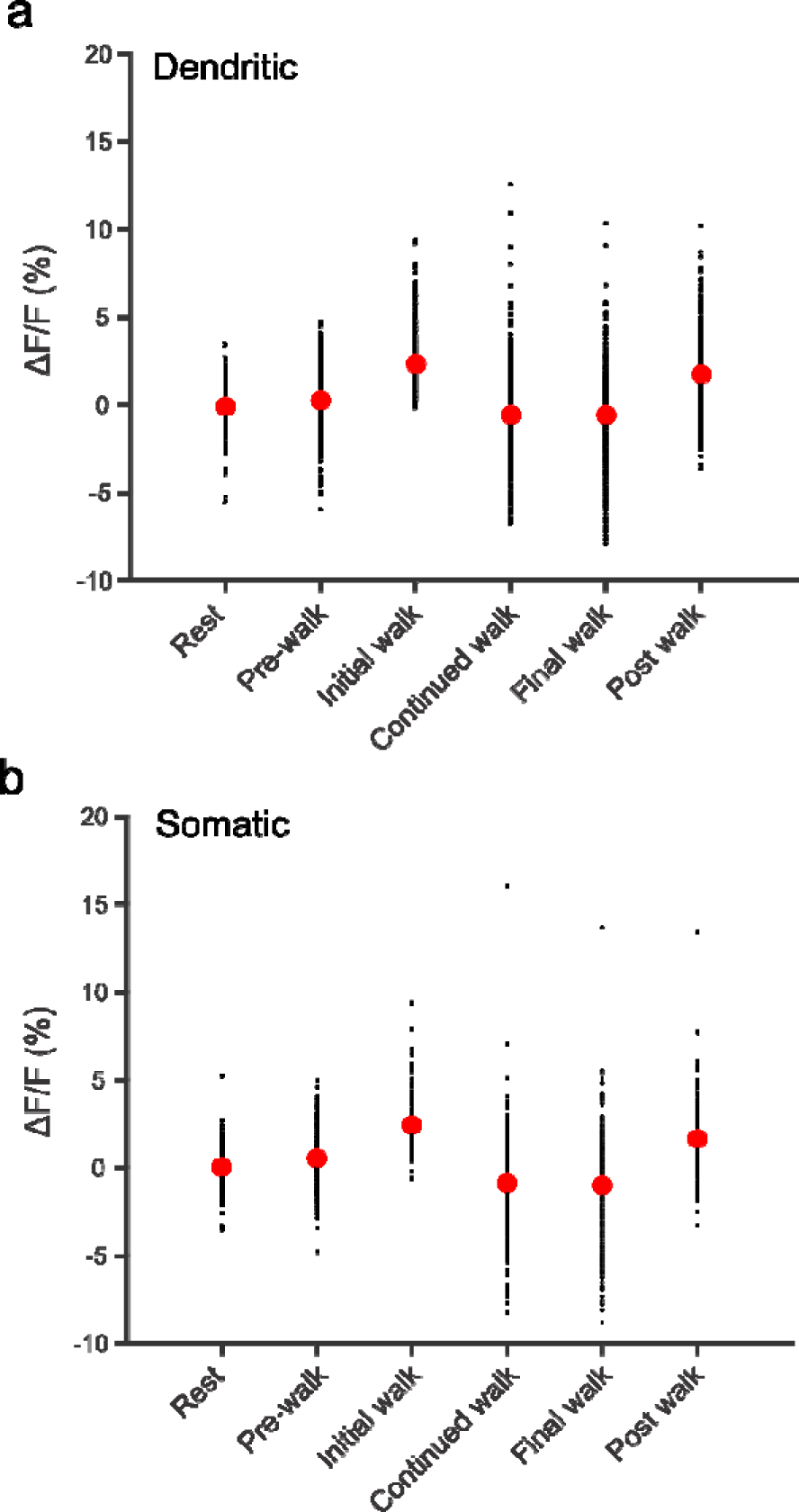
Population changes in fluorescence during locomotion epochs. a) Average ΔF/F of all dendritic ICs across each locomotion epoch. Red dots denote population mean. Locomotion epochs are associated with significant changes in activation (One-way ANOVA F(5,3270) = 231.82, p < 0.001), with significant increases beginning during the pre-walk epoch (0.25% ± 1.7 ΔF/F, p < 0.05, Tukey HSD), and the strongest increases observed during initial walking (2.3% ± 1.4 ΔF/F, p < 0.05, Tukey HSD). b) Same as for (a), but for somatic ICs. Similar to dendritic ICs, locomotion epochs are associated with significant changes in somatic activity levels (One-way ANOVA F(5,1698) = 136.31, p < 0.001), with significant increases beginning during the pre-walk epoch (0.5% ± 1.7 ΔF/F, p < 0.05, Tukey HSD), and the strongest increases observed during initial walking (2.4% ± 1.2 ΔF/F, p < 0.05, Tukey HSD).

**Extended data figure 5.**
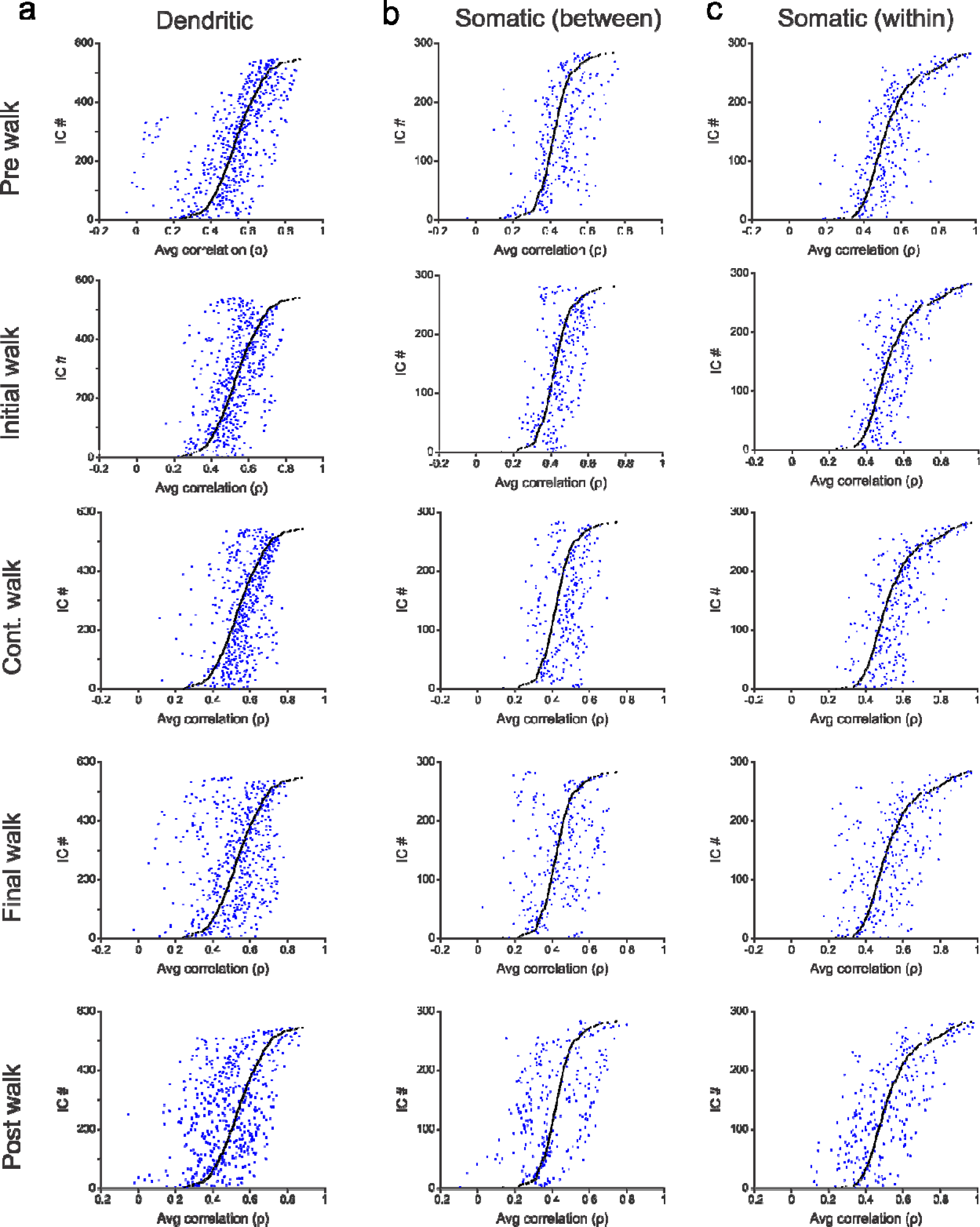
Changes in IC correlations during all walk periods organized by correlation at rest. a) Dot plots of mean IC domain correlations for dendritic ICs across all mice, sorted by correlation at rest (black dots). Blue dots denote the mean correlations for the sorted IC domains during pre-walk, initial walk, continued walk, final walk, and post walk period. b-c) Same as (a), but for somatic within and between IC correlations.

**Extended data figure 6.**
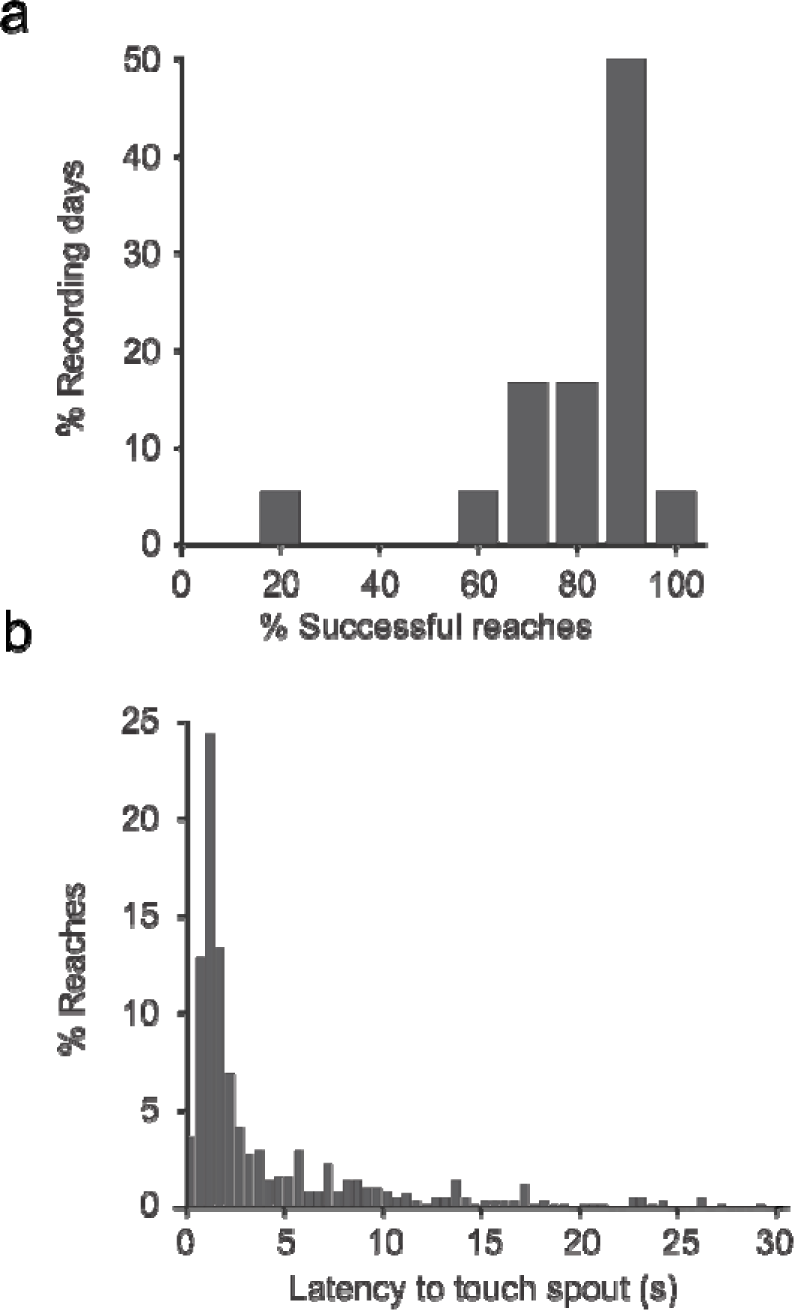
Characteristics of mouse reaching during standard reaching paradigm. a) Percent successful reaches across all recording days illustrate that mice reach at a high frequency, with > 50% of recording days showing a >90% successful reach rate. b) Distribution of the latency to touch the water spout for all reaches across all recording days. >50% of all reaches occur within 2 seconds of water droplet delivery. Only reaches that occurred within this 2 second window were used for subsequent analyses.

**Extended data figure 7.**
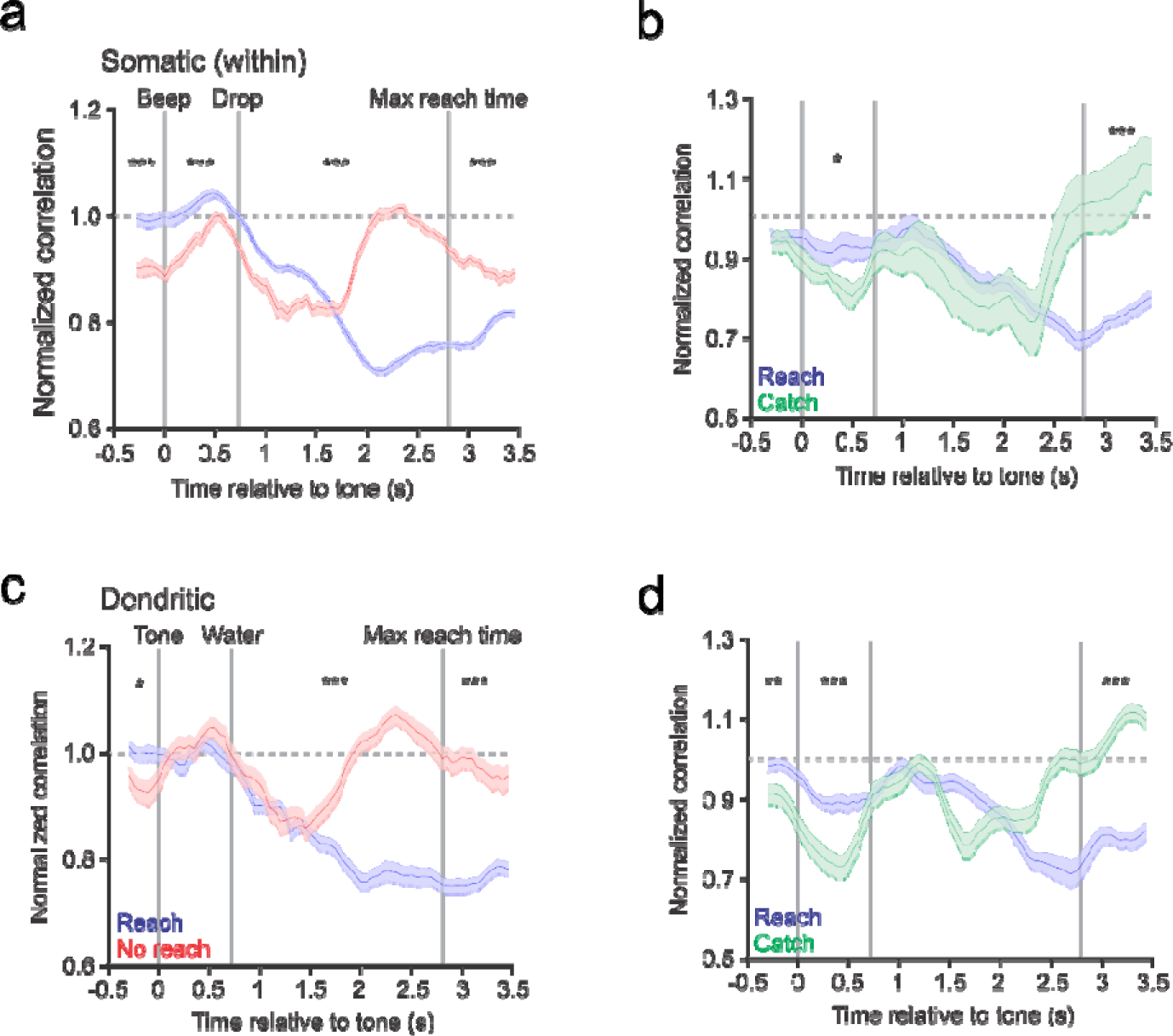
Somatic within IC and Dendritic between IC network dynamics during reaching. a-b) Average normalized correlations (relative to baseline, see Methods) for somatic IC domains within the same IC relative to auditory cue as for Fig. 7c-d. c-d) Same as for (a-b), but for dendritic IC domains. ***, p < 0.001; **, p < 0.01; *, p < 0.05, one-way ANOVAs.

## Methods

### Ethical approval

All experimental protocols were approved by the University of Minnesota’s Institutional Animal Care and Use Committee.

### Animals

For all experiments, mice were bred in-house and had *ad libitum* access to food and water in all housing conditions, except during the reaching paradigm, in which mice were water restricted to 1mL of water per day, in order to encourage reaching behavior. Mice expressing the Ca^2+^ indicator GCaMP6s selectively in Purkinje cells were generated by crossing mice expressing Cre selectively in Purkinje cells in the cerebellum (B6.Cg-Tg(Pcp2-cre)3555Jdhu/J; Jackson Laboratory stock 010536) with floxed-STOP GCaMP6s (B6.Cg-Igs7tm162.1(tetO-GCaMP6s,CAG-tTA2)Hze/J; Jackson Laboratory stock 031562). The resulting mice exhibited GCaMP6s expression that was selective to Purkinje cells, but in a sparse manner (**Extended data figure 1**), referred to as PcP2-GCaMP6s in the text, allowing for single cell resolution of Purkinje cells at the mesoscale level. Until implantation, animals were housed in standard housing conditions in the animal facility at the University of Minnesota. Following implantation, animals were singly housed in investigator managed housing on a 12-hour reverse light/dark cycle. All imaging experiments were performed during the dark (active) phase. A total of n = 8 mice of both sexes (n = 5 M, n = 3 F) were utilized for this study, with a minimum of n = 4 mice used for each behavioral paradigm.

### Cerebellar imaging window design and fabrication

Design and fabrication of the cerebellar imaging window and head fixation implants was adapted from previous procedures for imaging the cerebral cortex^25^. The curvature of the skull over the dorsal cerebellar cortex was mapped using a modified CNC mill, and a 2D profile of the skull surface was created to make the structural frame of the cerebellar window implant. A separate head fixation implant was created by modifying the previous cerebral cortical imaging windows (**Figure 1**). The window frame and head fixation implants were 3D-printed on a Form 2 printer (Formlabs) using Baclk v4 resin (RS-F2-GPBK, Formlabs). A small section of PET film was affixed to the window frame using epoxy, and bonded overnight in a custom, 3D printed mold. For the head fixation implant, three holes were tapped using an 0-80 hand tap for securing to a titanium headplate during the implantation surgery. Excess PET was carefully cut away from the frame and the final window cleaned with ethanol prior to implantation.

### Surgical procedures

Adult mice (postnatal day 45 or later) underwent implantation procedures under isofluorane anesthesia. Buprenorphine ER (1mg/kg) and Dexamethasone (2mg/kg) was administered subcutaneously for analgesia and inflammation, respectively. The scalp was removed to expose the skull over the dorsal surface of both the cerebral and cerebellar cortices, and the surface of the skull was scraped to remove the fascia. For window implantation, a large craniotomy matching the external profile of the cerebellar window frame was performed using a high speed drill, and the skull removed exposing lobules IV/V, VI, and VII of the cerebellar cortex, including the vermis, simplex, and portions of crus I and crus II. The window was placed over the cerebellar cortex and bonded to the skull using Vetbond (3M) and further secured using Metabond dental cement (C&B Metabond, Parkell Inc.). The head fixation implant was placed on the skull over the dorsal cerebral cortex and secured using Metabond. A custom titanium headplate was screwed onto the previously tapped holes in the head fixation implant, and the entire space between the window and headplate was filled with dental cement to protect the window from the external environment. A 3D-printed cap was attached to the titanium headplate to protect the PET surface. Post-operative care consisted of recovery from anesthesia on a heating pad with regular visual inspection, followed by daily post-operative monitoring for a minimum of three days to assess comfort level and healing of the surgical site. Animals were allowed to recover a minimum of five days prior to habituation to head fixation.

### Fluorescence imaging

Awake, mesoscale cerebellar imaging was performed under an epifluorescence microscope (Nikon AZ100) with a 2X objective. Using a digital zoom, the field of view was adjusted to image the dorsal cerebellum, with the focal plane adjusted to allow for resolution of both Purkinje cell dendrites and somata at a spatial resolution of 1024×1024 (total FOV ∼5 x 5mm). We found that this resolution and focal plane allowed for imaging somatic and dendritic activity throughout the entire cerebellar window (see **Figure 1**). Dual wavelength illumination was used to capture both Ca^2+-^-dependent (470 nm, blue light) and Ca^2+^-independent (405 nm, violet light) GCaMP6s signals on consecutive frames by a Cairn OptoLED driver (Cairn OptoLED, P1110/002/000; P1105/405/LED, P1105/470/LED). Ca^2+^-dependent and-independent signals were sampled by alternating the illumination of the LEDs, as described previously^26, 27, 55–57^. Single-photon fluorescence images were captured at a frequency of 20 fps, 18 ms exposure, using a high-speed CMOS camera (ORCA-Flash4.0 V3, Hamamatsu photonics) controlled with MetaMorph. Continuous images were collected as tiff stacks over 3.5-5.5 minute periods, which comprised a single imaging trial. A minimum of 4 wide-field stacks were collected for each recording day, and 3-10 recording days collected per mouse, per behavioral paradigm. For a subset of imaging days performed on the disk treadmill, both wide-field and higher magnification imaging stacks (n = 4 stacks for each magnification level) were obtained to characterize both mesoscale and single Purkinje cell ICs, respectively.

### Behavioral paradigms

The first set of behavioral experiments was performed with mice head-fixed on a freely moving disk treadmill to allow for spontaneous locomotion^26, 27^. Behavior was monitored and recorded using a high-speed IR-sensitive CMOS camera (Blackfly, FLIR systems) under infrared illumination that did not affect concurrent Ca^2+^ imaging. Synchronization of the behavior and microscope cameras was controlled by a series of TTL pulses delivered by Spike2 software and a CED power 1401 data acquisition system (Cambridge Electronic Design). To identify periods of locomotion, mouse forepaws and hindpaws were labeled using DeepLabCut for markerless behavioral tracking^33^, and the average broadband power for each paw was computed for a 1 s sliding window (50 ms step size) across each stack using custom Python code. Locomotion was reliably detected from this 1-20Hz average broadband power range (**Extended data figure 3**).

For cued the reaching experiments, the mice were water restricted following the protocol developed by Guo et al^58^. With the CMOS camera mounted from below, the underside of the mouse was recorded using a mirror, allowing for the resolution of limb position relative to the spout. Each trial of the reaching paradigm consisted of an auditory cue followed by the delivery of a water droplet through a water spout after a 700 ms delay. The water spout was connected to a capacitive sensor, to allow for the detection of the mouse’s paw making contact with the water spout. In a subset of imaging days, contact with the spout was recorded manually from the behavioral camera. The delivery of auditory tone and the water droplet, as well as monitoring of the capacitive sensor, was controlled using a Bpod finite state machine (Bpod, Sanworks) and custom written Matlab code. For each imaging stack, an initial 30 s baseline period was followed by 8 reaching trials, each separated by 30 s, resulting in 5.5 minute long stacks. A minimum of 4 stacks was collected per recording session, and 3-10 recording sessions collected per mouse.

For the catch trial experiments, stacks were equal in duration, with reach trials separated by the same 30 s window, but one of the auditory tones between trials 2-8 in each stack was randomly selected to not result in water delivery. As for the standard cued reaching paradigm, a minimum of 4 stacks was collected per recording session, and 3-10 recording sessions collected per mouse.

### Preprocessing pipeline

Preprocessing of Ca^2+^ data was performed for each recording day across all behavioral paradigms. First, fluorescence imaging stacks were deinterleaved based on the illumination wavelength (470 vs 405nm), and the first 10 s of imaging data removed to eliminate the initial rundown of the Ca^2+^ fluorescence signal. Ca^2+^-independent signals were removed from the 470nm data using a standard subtraction approach, as described previously^26, 27, 55–57^, and corrected for any small motion artifacts by co-registering each imaging frame to a reference image using a rigid transformation. The ΔF/F of each pixel was computed, and the resulting preprocessed imaging stacks were concatenated for a given recording day. Concatenated stacks were compressed using singular value decomposition (SVD), retaining the first 1000 principal components. Concatenated, SVD compressed data maintains important features of the original full-rank data while reducing the necessary computing time for subsequent analyses.

### Spatial independent component analysis

Spatial ICA was performed for each compressed dataset, for each mouse, computing 60 initial, unthresholded independent components (ICs) using the Joint Approximation and Diagonalization of Eigenmatrices (JADE) algorithm, as described previously^26^. Unthresholded spatial ICs were generated by multiplying each IC solution back into the original vector space and transforming into a z-score image. Binary masks of each IC were obtained by thresholding the pixels by both z-score and minimum domain size. First, all pixels of a given IC image were thresholded by z-score of ±3.5 or greater. Next, all groups of contiguous pixels throughout the thresholded image were selected. Each group of contiguous pixels with a size of 30 pixels or greater was determined to be an IC domain, such that each IC could possess multiple, non-contiguous domains. The resulting thresholded ICs were visually inspected for artifacts, such as areas overlying vasculature, that were manually discarded.

### Analysis of independent component domain modulation during locomotion

Broad characterization of Purkinje cell IC domain modulation during locomotion was first characterized by taking the average Δf/f for each IC during a behavioral epoch of interest. Locomotion behavior was first segmented by one of six epochs: rest, pre-walking (the 3 s prior to locomotion onset), initial walking (the first 3 s of locomotion), continued walking (the period of locomotion lasting from after the first 3 s of walking until the final 3 s of walking), final walking (the final 3 s prior to locomotion offset) and post walking (the first 3 s after locomotion offset). To avoid any overlap in locomotion epochs, periods of walking were thresholded such that they had to be a minimum of 3 seconds long, and any overlapping epochs (e.g. a post walk window for n walk bout that overlaps with the pre walk window for n + 1 walk bout) were discarded. For each IC domain, the average ΔF/F was computed across all time points in a given locomotion epoch, and the resulting mean ΔF/F was visualized across all IC domains using heat maps.

IC domain interactions were also quantified using correlation analyses. For each locomotion epoch, the IC domain ΔF/F was concatenated, and the resulting concatenated ΔF/F data was collected for all IC domains, and the correlation between each IC domain was computed. To quantify broad changes in correlations during locomotion, we computed the change in IC domain correlations compared to correlations at rest across all IC domains, all mice, and all recording days. Since somatic ICs consisted of numerous domains, correlations for IC domains that belonged to the same IC (i.e. “within IC”) and correlations for IC domains that belonged to different ICs (i.e. “between IC”) were also computed.

### Independent component domain spatial correlation analyses

To quantify the correlation between IC domains as a function of their spatial positioning, spatial correlation maps were computed for both dendritic domains (between IC) and somatic domains (within and between ICs). Correlations for all IC domains were collected as a function of their relative spatial positioning (x, y) over an 800 pixel, or approximately 4 x 4 mm region (**Figure 4A, cartoon schematics**). These spatial correlation maps were averaged across all IC domains, for all mice, and all recording days, resulting in a population spatial correlation map. The resulting maps were binned by a factor of 10, resulting in 80 x 80 pixel maps, over the same 4 x 4 mm spatial extent (**Figure 4A**). This spatial correlation analysis was performed for all locomotion epochs, and changes in spatial correlations for a locomotion epoch of interest was computed as the difference between the population spatial correlation map at rest versus the locomotion epoch of interest (**Figure 4C**).

### Analysis of IC domain modulation during reaching

For the cued reaching paradigm, all reaches were first thresholded so that the analysis was performed only on trials in which the mouse touched the spout within two seconds of water delivery. For these successful reaches, IC domain modulation was quantified by aligning IC domain ΔF/F to a behavioral event of interest (auditory tone, water droplet delivery, or spout touch). The resulting ΔF/F was binned into temporal windows, averaged, and plotted as heat maps as for locomotion analyses, resulting in ΔF/F maps at time points relative to each behavioral event. IC domain spatial correlation maps were computed as for locomotion, with the rest correlation computed from the first 20 s of a given trial, in which no auditory tone is delivered. Note that this 20 s period is followed by the initial 30 s waiting period prior to the first auditory tone delivery, so the mouse should not be anticipating an auditory tone during this time. The changes in spatial correlation were computed by subtracting the population spatial correlation at rest from either the population spatial correlations during the tone period (the time of auditory tone up until the time of water droplet delivery), or the population spatial correlations during the reach period (the time of water delivery up until the time at which the mouse touches the spout).

Similar analyses were performed for the catch trial paradigm for both average ΔF/F of IC domains and changes in IC domain spatial correlations, with the change in correlation maps being determined by the difference in spatial correlation during the standard reach condition (tone followed by water delivery) from the spatial correlation during the catch condition (tone followed by no water delivery).

### Population modulation maps during reaching

In order to quantify the overall modulation of IC domains across the population, the ΔF/F of each IC domain was aligned to the time of auditory tone delivery in three different conditions: trials in which the auditory tone and water delivery was followed by a successful reach within 2 s of water (“reach”), trials in which the auditory tone and water was not followed by a reach (“no reach”), and trials from the catch trial paradigm, in which the tone was not followed by water (“catch”). The ΔF/F for each time point relative to the auditory tone was averaged into 100 ms bins. For each time bin, the IC domain modulation was converted into a z-score by comparing the average ΔF/F to the ΔF/F during the rest period (the first initial 20 s of a recording stack, as for the spatial correlation analyses). The collected z-score time courses were collected for all IC domains across all mice, all recording days, and thresholded by a minimum z-score of ±2.5. These maps were then averaged, resulting in an overall quantification of significant modulation across the population for each time bin relative to the auditory tone, for each behavioral outcome condition.

### Sliding window correlation analyses

To quantify changes in correlated IC domain activity across the population, IC domain ΔF/F was aligned to the auditory tone in each behavioral outcome condition as above. Next, for each IC domain, we computed the correlations between a given IC domain and all other IC domains in a 1 s sliding window, beginning 300 ms prior to the delivery of the auditory tone. These temporal IC domain correlations were normalized to the correlation at rest, resulting in a timecourse of the change in correlation over time for each behavioral outcome.

### Statistical analyses

Significant changes in correlations between somatic and dendritic ICs as a function of relative spatial distance, and changes in correlations for mediolateral versus anteroposterior dimensions were determined using t-tests followed by a the Benjimini and Hostberg^54^ false discovery rate correction (p < 0.05). Broad changes in spatial correlation maps, including during pre-walk and post-walk periods, as well as proximal versus distal changes during reach epochs, were determined using one-way ANOVAs and (where applicable) post-hoc Tukey HSD tests on the collected change in spatial correlation map bins of interest (p < 0.05). For the sliding window analysis, significant changes in correlations in different behavioral conditions were determined from the average normalized correlation during each behavioral window of interest (pre-tone: -300:0 ms, tone: 0:700 ms; reach: 700:2700 ms; post-reach: 2700-3500 ms) for each IC domain using one-way ANOVAs followed by post-hoc Tukey HSD tests (p < 0.05).

## Notes

### Competing Interest Statement

The authors have declared no competing interest.

## Reference

1. Eccles, J.C., Ito, M. & Szentágothai, J.n. The cerebellum as a neuronal machine (Springer-Verlag, Berlin, New York etc., 1967).

2. Itō, M. The cerebellum and neural control (Raven Press, New York, 1984).

3. Braitenberg, V. Functional interpretation of cerebellar histology. Nature 190, 539–540 (1961).

4. Cramer, S.W., Gao, W., Chen, G. & Ebner, T.J. Reevaluation of the beam and radial hypotheses of parallel fiber action in the cerebellar cortex. J Neurosci 33, 11412–11424 (2013).

5. Ozol, K., Hayden, J.M., Oberdick, J. & Hawkes, R. Transverse zones in the vermis of the mouse cerebellum. J Comp Neurol 412, 95–111 (1999).

6. Voogd, J. & Glickstein, M. The anatomy of the cerebellum. Trends Cogn Sci 2, 307–313 (1998).

7. Brodal, A. & Kawamura, K. Olivocerebellar projection: a review. Adv Anat Embryol Cell Biol 64, IVIII, 1–140 (1980).

8. Reeber, S.L., White, J.J., George-Jones, N.A. & Sillitoe, R.V. Architecture and development of olivocerebellar circuit topography. Front Neural Circuits 6, 115 (2012).

9. Schmahmann, J.D. From movement to thought: anatomic substrates of the cerebellar contribution to cognitive processing. Hum Brain Mapp 4, 174–198 (1996).

10. Zhou, H., et al. Cerebellar modules operate at different frequencies. Elife 3, e02536 (2014).

11. Beckinghausen, J. & Sillitoe, R.V. Insights into cerebellar development and connectivity. Neuroscience Letters 688, 2–13 (2019).

12. Apps, R., et al. Cerebellar Modules and Their Role as Operational Cerebellar Processing Units: A Consensus paper [corrected]. Cerebellum 17, 654–682 (2018).

13. Cerminara, N.L., Lang, E.J., Sillitoe, R.V. & Apps, R. Redefining the cerebellar cortex as an assembly of non-uniform Purkinje cell microcircuits. Nat Rev Neurosci 16, 79–93 (2015).

14. De Zeeuw, C.I. Bidirectional learning in upbound and downbound microzones of the cerebellum. Nat Rev Neurosci 22, 92–110 (2021).

15. De Zeeuw, C.I., Lisberger, S.G. & Raymond, J.L. Diversity and dynamism in the cerebellum. Nat Neurosci 24, 160–167 (2021).

16. Manni, E. & Petrosini, L. A century of cerebellar somatotopy: a debated representation. Nat Rev Neurosci 5, 241–249 (2004).

17. Apps, R. & Hawkes, R. Cerebellar cortical organization: a one-map hypothesis. Nat Rev Neurosci 10, 670–681 (2009).

18. Lang, E.J., Sugihara, I., Welsh, J.P. & Llinas, R. Patterns of spontaneous purkinje cell complex spike activity in the awake rat. J Neurosci 19, 2728–2739 (1999).

19. Sasaki, K., Bower, J.M. & Llinas, R. Multiple Purkinje Cell Recording in Rodent Cerebellar Cortex. Eur J Neurosci 1, 572–586 (1989).

20. Welsh, J.P., Lang, E.J., Suglhara, I. & Llinas, R. Dynamic organization of motor control within the olivocerebellar system. Nature 374, 453–457 (1995).

21. Heffley, W., et al. Coordinated cerebellar climbing fiber activity signals learned sensorimotor predictions. Nat Neurosci 21, 1431–1441 (2018).

22. Kostadinov, D., Beau, M., Blanco-Pozo, M. & Hausser, M. Predictive and reactive reward signals conveyed by climbing fiber inputs to cerebellar Purkinje cells. Nat Neurosci 22, 950–962 (2019).

23. Wagner, M.J., Kim, T.H., Savall, J., Schnitzer, M.J. & Luo, L. Cerebellar granule cells encode the expectation of reward. Nature 544, 96–100 (2017).

24. Wagner, M.J., et al. A neural circuit state change underlying skilled movements. Cell 184, 3731–3747 e3721 (2021).

25. Ghanbari, L., et al. Cortex-wide neural interfacing via transparent polymer skulls. Nat Commun 10, 1500 (2019).

26. Nietz, A.K., et al. To be and not to be: wide-field Ca2+ imaging reveals neocortical functional segmentation combines stability and flexibility. Cereb Cortex 33, 6543–6558 (2023).

27. West, S.L., et al. Wide-Field Calcium Imaging of Dynamic Cortical Networks during Locomotion. Cereb Cortex (2021).

28. Ramirez, J.E. & Stell, B.M. Calcium Imaging Reveals Coordinated Simple Spike Pauses in Populations of Cerebellar Purkinje Cells. Cell Rep 17, 3125–3132 (2016).

29. Makino, H., et al. Transformation of Cortex-wide Emergent Properties during Motor Learning. Neuron 94, 880–890 e888 (2017).

30. Michikawa, T., et al. Distributed sensory coding by cerebellar complex spikes in units of cortical segments. Cell Rep 37, 109966 (2021).

31. Ruigrok, T.J. Ins and outs of cerebellar modules. Cerebellum 10, 464–474 (2011).

32. Voogd, J. Cerebellar zones: a personal history. Cerebellum 10, 334–350 (2011).

33. Mathis, A., et al. DeepLabCut: markerless pose estimation of user-defined body parts with deep learning. Nat Neurosci 21, 1281–1289 (2018).

34. Muzzu, T., Mitolo, S., Gava, G.P. & Schultz, S.R. Encoding of locomotion kinematics in the mouse cerebellum. PLoS One 13, e0203900 (2018).

35. Sauerbrei, B.A., Lubenov, E.V. & Siapas, A.G. Structured Variability in Purkinje Cell Activity during Locomotion. Neuron 87, 840–852 (2015).

36. Udo, M., Matsukawa, K., Kamei, H., Minoda, K. & Oda, Y. Simple and complex spike activities of Purkinje cells during locomotion in the cerebellar vermal zones of decerebrate cats. Exp Brain Res 41, 292–300 (1981).

37. Schweighofer, N., Doya, K. & Lay, F. Unsupervised learning of granule cell sparse codes enhances cerebellar adaptive control. Neuroscience 103, 35–50 (2001).

38. Galliano, E., et al. Silencing the majority of cerebellar granule cells uncovers their essential role in motor learning and consolidation. Cell Rep 3, 1239–1251 (2013).

39. Albus, J.S. A theory of cerebellar function. Mathematical Biosciences 10, 25–61 (1971).

40. Marr, D. A theory of cerebellar cortex. J Physiol 202, 437–470 (1969).

41. Quarta, E., et al. Distributed and Localized Dynamics Emerge in the Mouse Neocortex during Reach-to-Grasp Behavior. J Neurosci 42, 777–788 (2022).

42. Findling, C., et al. Brain-wide representations of prior information in mouse decision-making. bioRxiv, 2023.2007.2004.547684 (2023).

43. Laboratory, I.B., et al. A Brain-Wide Map of Neural Activity during Complex Behaviour. bioRxiv, 2023.2007.2004.547681 (2023).

44. Steinmetz, N.A., et al. Neuropixels 2.0: A miniaturized high-density probe for stable, long-term brain recordings. Science 372 (2021).

45. Steinmetz, N.A., Zatka-Haas, P., Carandini, M. & Harris, K.D. Distributed coding of choice, action and engagement across the mouse brain. Nature 576, 266–273 (2019).

46. Person, A.L. & Raman, I.M. Purkinje neuron synchrony elicits time-locked spiking in the cerebellar nuclei. Nature 481, 502–505 (2011).

47. Gauck, V. & Jaeger, D. The control of rate and timing of spikes in the deep cerebellar nuclei by inhibition. J Neurosci 20, 3006–3016 (2000).

48. Sedaghat-Nejad, E., Pi, J.S., Hage, P., Fakharian, M.A. & Shadmehr, R. Synchronous spiking of cerebellar Purkinje cells during control of movements. Proc Natl Acad Sci U S A 119, e2118954119 (2022).

49. Herzfeld, D.J., Joshua, M. & Lisberger, S.G. Rate versus synchrony codes for cerebellar control of motor behavior. Neuron 111, 2448–2460 e2446 (2023).

50. Person, A.L. & Raman, I.M. Synchrony and neural coding in cerebellar circuits. Front Neural Circuits 6, 97 (2012).

51. Nashef, A., Spindle, M.S., Calame, D.J. & Person, A.L. A dual Purkinje cell rate and synchrony code sculpts reach kinematics. bioRxiv (2023).

52. Heck, D.H., De Zeeuw, C.I., Jaeger, D., Khodakhah, K. & Person, A.L. The neuronal code(s) of the cerebellum. J Neurosci 33, 17603–17609 (2013).

53. Chen, T.W., et al. Ultrasensitive fluorescent proteins for imaging neuronal activity. Nature 499, 295–300 (2013).

54. Benjamini, Y. & Hochberg, Y. Controlling the False Discovery Rate: A Practical and Powerful Approach to Multiple Testing. Journal of the Royal Statistical Society: Series B (Methodological) 57, 289–300 (1995).

55. Allen, W.E., et al. Global Representations of Goal-Directed Behavior in Distinct Cell Types of Mouse Neocortex. Neuron 94, 891–907 e896 (2017).

56. Ren, C. & Komiyama, T. Characterizing Cortex-Wide Dynamics with Wide-Field Calcium Imaging. J Neurosci 41, 4160–4168 (2021).

57. Couto, J., et al. Chronic, cortex-wide imaging of specific cell populations during behavior. Nat Protoc 16, 3241–3263 (2021).

58. Guo, Z.V., et al. Procedures for behavioral experiments in head-fixed mice. PLoS One 9, e88678 (2014).

